# Population genomics for coral reef restoration - a case study of staghorn corals in Micronesia

**DOI:** 10.1101/2024.09.29.615720

**Authors:** Dareon Rios, Hector Torrado, Sarah Lemer, Crawford Drury, David Burdick, Laurie Raymundo, David J. Combosch

## Abstract

Staghorn *Acropora* corals are ecological keystone species in shallow lagoons and back reef habitats throughout the tropics. Their widespread decline coupled with their amenability for asexual propagation propelled them to the forefront of global coral restoration efforts - albeit frequently without much scientific input. To guide these efforts and as a blueprint for similar projects, we conducted a comprehensive population genomic study of *Acropora cf. pulchra*, a major restoration target species in the Indo-Pacific. Our results revealed that *A.* cf. *pulchra* populations in the Mariana Islands are characterized by large clonal clusters and extremely low levels of genetic diversity. Differentiation among populations followed a significant isolation-by-distance pattern and delineated two distinct metapopulations on Guam. Our investigation identified critical population genetic parameters, necessitating targeted management strategies, and provides actionable guidelines for effective conservation efforts. For management and conservation, two populations emerged as pivotal connectivity hubs with elevated genetic diversity. For restoration, we show that *A*. cf. *pulchra* populations demonstrated a suitability for extensive asexual propagation and provide guidelines how to best apply that. To preserve and augment genetic diversity, strategies to mitigate inbreeding are crucial until sexual reproduction can be fully integrated into restoration protocols. Critical sites for restoration include local connectivity hubs, fringing lagoons that connect metapopulations, and back reefs around a particularly isolated population. These findings offer crucial insights into the genetic landscape of a keystone coral species and provide actionable recommendations for coral conservation and restoration. By advocating for the preservation of population connectivity and the promotion of genotypic, genetic, and symbiont diversity in coral restoration, our study serves as a blueprint for leveraging population genomic studies to enhance the efficacy and resilience of restoration projects on remote islands.

## 1 INTRODUCTION

Coral reefs are declining rapidly worldwide due to increasing seawater temperature, ocean acidification, and local anthropogenic stressors (Hoegh-Guldberg et al. 2007, 2018, Zande et al. 2019). Over the past three decades, reef-building corals have faced huge losses, with one-third of reef corals being at risk of extinction (Carpenter et al. 2008, Mumby and Steneck 2008). Globally, coral reefs play vital functional roles contributing to economic growth, serving as coastal protection, providing habitats for various marine species, and sustaining cultural and traditional practices (Hicks 2011). Reef managers have therefore turned to coral restoration as one important tool for the management and preservation of tropical coral reefs.

Scleractinian corals are widespread and form most of the framework of modern coral reefs. *Acropora*, the most abundant coral genus with around 149 species (Cowman et al. 2020, Ball et al. 2022), is most diverse in the coral triangle, where it provides much of the reef structure, supporting the most diverse marine life (Wallace et al. 1999, Veron and Stafford-Smith 2000). While *Acropora* corals are hermaphroditic broadcast spawners, their local abundance often depends on asexual fragmentation (Tunnicliffe 1981, Highsmith 1982), which enables rapid propagation but also increases vulnerability (e.g. Bruckner 2002, Vollmer and Palumbi 2007, Drury et al. 2016, 2017). This is particularly true for staghorn *Acropora* corals, a fast-growing group that dominates sheltered areas and relies heavily on vegetative fragmentation. Staghorns can recover locally but require successful fertilization and dispersal for distant recolonization, which is often limited (e.g. (Highsmith 1982, Drury et al. 2016, 2017) but see (Gilmour et al. 2013))

On Guam, staghorn *Acropora* are locally-dominant reef-builders that form vital habitats for local fishes and invertebrates on shallow reef flats and lagoonal patch reefs (Raymundo et al. 2017). Guam’s staghorn *Acropora* have been impacted by various stressors including infectious disease (Myers and Raymundo 2009), *Drupella* and *Acanthaster planci* predation and outbreaks (Burdick et al. 2008), and widespread coral bleaching (Raymundo et al. 2017, 2019). Most notably, staghorn populations suffered an estimated 50% loss in coral cover over a three-year period (2013– 2015), marked by consecutive bleaching events and extreme low tides (Reynolds et al. 2014, Raymundo et al. 2017, 2019). On Saipan, 200 km north of Guam, a >90% loss of staghorn *Acropora* spp. was observed in the main lagoon during these bleaching events (BECQ-DCRM, Long-Term Monitoring Program, unpublished data). In response to this recent and drastic decline, and because of their suitability for extensive asexual propagation, staghorn *Acropora* corals are a major restoration target, on Guam and worldwide (Boström-Einarsson et al. 2020), and *A. pulchra* (Brook, 1891) is one of the main target species throughout the Indo-West Pacific (e.g. (Soong and Chen 2003, Borell et al. 2010, Cruz et al. 2014, Romatzki 2014, DeMars 2021, Raymundo et al. 2022).

Conserving existing biodiversity takes precedence over restoration, but when not all populations can be protected, informed trade-offs are necessary. Populations can be prioritized based on genetic and adaptive diversity (DeWoody et al. 2021, Teixeira and Huber 2021, Willi et al. 2022) or their role in connectivity (Jones et al. 2007, Hoban 2018, Beger et al. 2022, Fontoura et al. 2022). Population genetics is essential for conservation, as it uncovers key evolutionary patterns and processes shaping species presence and distribution (e.g. (Vellend and Geber 2005, Allendorf et al. 2012, Breed et al. 2019). In addition, population genetics connects evolutionary and ecological processes that are crucial in aiding management efforts for sustaining reef biodiversity and functioning and provides critical basic knowledge about the restoration targets (Vellend and Geber 2005, Falk et al. 2006, Richards et al. 2016, Breed et al. 2019).

Here, we assess the genetic composition of one of the main coral restoration target species in the Indo-Pacific, *A.* cf. *pulchra* to guide management and restoration. Although the species life history and reproduction (e.g. (Harrison et al. 1984, Babcock et al. 2003, Baird et al. 2009, Darling et al. 2012, Lapacek 2017), general ecology (e.g. (Veron 1986, Díaz and Madin 2011, Muir et al. 2015) and major symbionts (e.g. (Li et al. 2008, Edmunds et al. 2014) are well known, important open questions concern its systematics, taxonomy and heat tolerance (Cowman et al., Reuter et al., in prep) as well as its population genetics (Hein et al. 2021, Vardi et al. 2021, Shaver et al. 2022, Suggett et al. 2024), which is the focus of the present study. Our goal was to conduct a comprehensive population genomic assessment as a blueprint for conservation and restoration genomic studies elsewhere. We specifically assessed the following vital aspects to evaluate their importance and suitability for informing management and conservation in small island states that are particularly challenged by global climate change (Hernández-Delgado 2024):

a. The extent of clonality and the spatial distribution of clones within and among populations, which provides a baseline of genotypic diversity in wild populations to assess the suitability of asexual propagation and help to decide where and how fragments for propagation should be harvested and replanted to efficiently maximize genotypic diversity (e.g. (Reynolds et al. 2012, Koch 2021, Nef et al. 2021).
b. The genetic diversity of the target species, to assess its evolutionary potential (O’Grady et al. 2004, Kardos et al. 2021), adaptive capacity (e.g. (Haig 1998, Reed and Frankham 2003, Oppen and Gates 2006, DiBattista 2008, Shearer et al. 2009) and the need for intervention and active restoration (e.g. (Spielman et al. 2004, Frankham et al. 2013).
c. The population structure, migration and distribution of related individuals among populations, which can identify potential barriers to connectivity and important source populations for conservation and restoration (e.g. (Palumbi 2003, Leiva et al. 2022, Shaver et al. 2022).
d. Signatures of selection, which informs on the extent of localized adaptations and what environmental factors might be driving localized adaptations, to extrapolate findings beyond surveyed populations (e.g. (Mijnsbrugge et al. 2010).
e. Dominant symbionts genera, to assess how specific the relationship between the host and its primary symbionts is and map the spatial distribution of dominant symbionts, which may provide additional opportunities to harden holobionts to further environmental change (e.g. (Dixon et al. 2015, Anthony et al. 2017, Morikawa and Palumbi 2019, Schoepf et al. 2019).

Specifically, we analyzed genome-wide ddRADseq data for 188 *A.* cf. *pulchra* samples to quantify patterns of population genetics within and among populations around Guam and between Guam and Saipan, the two main islands of the Marianas, Micronesia.

## 2 METHODS

### 2.1 Sampling sites and process

*Acropora* cf. *pulchra* samples were collected between May 2018 and October 2019 from five locations around Guam (Fig. 1, Table S1). Populations were selected to maximize geographic distances and environmental differences between sites. For example, Urunao and Togcha occur close to the reef crest on shallow (<0.5 m) reef flat platforms, while populations in Agat, Cocos, and West Agaña are located in wider and deeper (∼1+ m) back reef lagoons at greater distance from the reef crest. In addition, forty-one staghorn *Acropora* samples collected across Saipan Lagoon. Out of these 267 staghorn *Acropora* samples, 233 were identified as *A.* cf. *pulchra* (see below) and 188 yielded sufficient sequencing data to be analyzed in detail (Table S1).

**FIGURE 1:** Location of the five sampling sites of *A.* cf. *pulchra* on Guam and Saipan. **Submitted as Figure1_20231211.pdf**

Samples were collected at depths between 0.5 and 1 m, every 10 meters along transects to minimize the collection of clonemates and assess small scale spatial genetic structures. In Togcha, the limited spatial extent of the local staghorn *Acropora* population required random sampling and the samples from Saipan were collected haphazardly as well. Underwater photographs were taken of each sampled colony and small nubbin samples were carefully removed with a wire cutter, placed in falcon tubes filled with seawater and transported back to the University of Guam (UOG) Marine Laboratory. Upon arrival, tissue samples were preserved in 95% ethanol and stored in a - 20C freezer. Remaining nubbins were bleached and cataloged in the UOG Biorepository as skeletal vouchers (catalog numbers #130-199, #519-674).

### 2.2 DNA extraction and ddRAD library preparation

Total genomic DNA was extracted using the DNAeasy Kit (Qiagen, Hildesheim, Germany) and the GenCatch Genomic DNA Extraction Kit (Epoch, Sugar Land, TX) following optimized manufacturer’s protocols. DNA quantity was measured with a Qubit 3.0 dsDNA fluorometer (Thermo Fisher Scientific Inc., Waltham, MA).

Double-digest restriction site-associated DNA (ddRAD) libraries were prepared in-house (Combosch et al. 2017)following a modified protocol based on (Peterson et al. 2012) and (Combosch et al. 2017). In brief, extracted DNA was digested using two high-fidelity restriction enzymes, NsiI and MspI. Resulting fragments were ligated to custom P1 and P2 adaptors with sample-specific barcodes and primer annealing sites. Barcoded samples were pooled into libraries, and size-selected (320-420 bp) with an E-Gel Size Select II Agarose Gel (Thermo Fisher Scientific Inc., Waltham, MA). Size-selected fragments were PCR-amplified using a high-fidelity polymerase (New England Biolabs, Ipswich, MA) with primers containing additional indices and flow cell annealing sites. Between 2 and 10 individual PCR reactions were set up per library and pooled subsequently to increase the diversity of sequencing pools. Between 15 and 22 PCR cycles (95°C for 30 s, 65°C for 30 s, 72°C for 60 s, with an initial denaturation step at 98°C for 30 s, and a final extension step at 72°C for 5 min) were used, depending on the concentrations of the resulting libraries.

Libraries were cleaned to remove excess adapters and primers using Agilent beads (Agilent Technologies, Santa Clara, CA) at a 1:0.6 library to beads ratio. Quality and quantity checks were performed on an Agilent Bioanalyzer 2100 (Agilent Technologies, Santa Clara, CA) and Qubit 3.0 (Thermo Fisher Scientific Inc., Waltham, MA), respectively. Lastly, libraries were single-end sequenced (120 bp) on an Illumina NextSeq500 Illumina (New England Biolabs, Ipswich, MA) at the University of Guam Marine Laboratory. Nine random samples were sequenced twice and analyzed separately, as 18 technical replicates, as well as combined in subsequent analyses.

### 2.3 Data curation and genotyping

Corals are clonal organisms with complex phylogenies, so data was analyzed in hierarchical approach that included a) identification and removal of cryptic species using phylogenomics, b) identification of clonemates for genotypic diversity analyses, followed by c) detailed population genomics analyses. Genomics datasets were processed using genotype likelihoods and hard-calling genotypes to accommodate different downstream software.

Raw reads were quality-trimmed with TrimGalore 0.6.5 (Krueger et al. 2023) with default settings to remove reads with an average quality score <30 and shorter than 36 bp. Resulting reads were demultiplexed using a custom python script (identify_dbrs6.py, H. Weigand, personal communication) to remove reads with uncalled bases, incomplete barcodes or restriction cut sites and trimmed to 100bp to remove lower quality sites. Cleaned and trimmed reads were aligned to the closely related *A. millepora* genome (Ying et al. 2019); Torrado et al. 2024) using Bowtie2 v2.3.5 (Langmead and Salzberg 2012), with default settings but excluding soft matches. Aligned reads were converted to bam files and sorted using SAMtools (Li et al. 2009).

Genotyping was performed using two separate approaches: ANGSD v0.93 (Korneliussen et al. 2014) and STACKS v2.3 (Rochette et al. 2019). For a subset of analyses that do not accommodate genotype probabilities, STACKS v2.3 was used to generate fixed genotype calls. STACKS was used in the reference-based mode and single nucleotide polymorphisms (SNPs) were identified using a Bayesian model with an alpha threshold of 0.05 for discovering SNPs (Catchen et al. 2011, 2013, Rochette et al. 2019). The STACKS populations program was then used to retain only loci that were present in 50% of all samples. This phylogenomic dataset was used for phylogenomic analyses to ensure only *A.* cf. *pulchra* samples were used in population genetic analyses. A preliminary phylogenomic analysis of closely related *Acropora* species was conducted with RaxML version 8.2.12 (Stamatakis 2014) on the CIPRES web portal (Miller et al. 2010), using the GTR model of sequence evolution with free model parameters estimated by RAxML (Fig. S1). Based on this tree, 233 *A.* cf. *pulchra* samples were identified as *A.* cf *pulchra* among the 267 genotyped *Acropora* samples (Table S1). Subsequent population genetic analyses (e.g. Fig. S2) did not indicate any significant outlier samples, tentatively confirming this approach.

For population genomic analyses, only unique (i.e. non-clonal) *A.* cf. *pulchra* samples with more than 5,000 high-quality mapped raw reads were used (n = 170, Table S1). For population genetic summary statistics, a dataset with all positions was used to avoid sample size biases (Schmidt et al. 2021) and a lower alpha threshold for discovering SNPs (0.01 instead of 0.05, as recommended in the STACKS manual) since the strict VCF filtering described below was not possible for datasets that include monomorphic positions. Other population genomic analyses included only the first SNP per ddRAD locus to avoid linkage between SNPs. VCFtools v1.13 was then used to remove individual variants with a coverage below 5x, loci absent in more than 50% of all samples, and loci with a coverage higher than 1.5 times the interquartile range of the dataset. Finally, loci with a major allele frequency equal or higher than 0.95 (i.e., virtually monomorphic) were identified using the ‘isPoly’ function in the ‘adegenet’ (Jombart 2012) R package and removed with VCFtools. This filtered dataset was used for population differentiation indices (FST, GST and DEST), AMOVAs, migration analyses (BA3-SNP) and the selection analyses with BayPass and BayesScan.

ANGSD generates genotype probabilities instead of fixed genotype calls. This approach incorporates genotype uncertainty, which is useful for low and variable coverage data (Korneliussen et al. 2014). ANGSD was run using the following filters: minimum mapping quality score of 20, minimum base call quality of 30, a minimum allele frequency of 0.05, a polymorphism threshold of 2 x 10^-6^, genotyped in at least 50% of samples, and a filter that retained only uniquely mapped reads. This full-locus ANGSD genotype likelihood dataset was used to calculate identity-by-state (IBS) and Thetastat. For other population genomic analyses, a filtered ANGSD dataset was generated by exporting genotype likelihoods as bcf files and removing all but 1 SNP per ddRAD locus using VCFtools v1.13. The same vcf filters for coverage and presence/absence as described above were applied to this dataset as well. As before, loci with a major allele frequency equal or higher than 0.95 (i.e., virtually monomorphic) were identified using the ‘isPoly’ function on ‘adegenet’ (Jombart 2012) R package and removed with VCFtools. This filtered VCF was used as input for ANGSD to perform the principal coordinates analysis (PCoA), ngsAdmix and ngsRelate.

### 2.4 Intra-population genomics: Clones, Relatedness and Spatial Genetic Structure (SGS)

To examine clonality, ANGSD was used to generate an identity-by-state (IBS) matrix following (Manzello et al. 2019) and (Barfield et al. 2018) using the R function hclust() and the method “average”. To determine a genetic distance threshold for identifying clones, a binned gap analysis (Fig. S2) was used to compare levels of relatedness between almost identical clonemates and unique genotypes. Technical replicates were used to determine a lower threshold. Results were displayed on a hierarchical clustering dendrogram with branch lengths displaying levels of genetic similarity (Fig. S2). Samples that exhibited lower genetic distances than the clonality threshold were identified as clones (Fig. S2). Clonality per population was calculated as the proportion of unique genotypes (NG/N), i.e. the genet/ramet ratio and denotes relative genotypic diversity. Genotypic evenness, indicating how evenly genotypes are present within populations, was calculated as the evenness of the effective number of genotypes across populations using GenoDive (Meirmans and Tienderen 2004). Subsequently, clonal genotypes were pruned to leave only a single representative with the highest number of mapped reads from each genet for downstream population genetic analyses.

To further investigate the relatedness among samples, the ANGSD subprogram NgsRelate was used to calculate pairwise relatedness (Rab; (Hedrick and Lacy 2015, Hanghøj et al. 2019) based on genotype likelihoods and population allele frequencies (Korneliussen and Moltke 2015). The average relatedness (Rab) measures the proportion of homologous alleles shared by 2 individuals, which is ∼0.5 between first-degree relatives, ∼0.25 between second-degree relatives, and ∼0.125 between third-degree relatives. Pairwise relationships were therefore binned as follows:

0.09375 - 0.1875 = Third-degree relatives, e.g. first cousins or great grandparents-grandchildren.

0.1875 - 0.375 = Second-degree relatives, e.g. aunts/uncles-nieces or grandparents-grandchildren.

>0.375 = First-degree relatives, e.g. parent-child or full siblings.

Fine-scale Spatial Genetic Structure (SGS) was estimated using the program SPAGeDi 1.5 (Hardy and VEKEMANS 2002). Loiselle’s kinship coefficient (Loiselle et al. 1995) was calculated over all samples within 10m intervals up to 200m for both the ramet dataset, i.e. including clones, and a genet only dataset, excluding clones. The 95% confidence intervals and standard errors were estimated based on 10,000 permutations of the genetic and the spatial datasets. Kinship values outside the 95% confidence intervals were interpreted as significant SGS at the spatial distance. The Sp statistic (Vekemans and Hardy 2004) was calculated using the rSpagedi function SpSummary (Browne 2019). The genetic patch size is the distance that corresponds to the first x-intercept of the kinship correlogram (Verity and Nichols 2014). Error bars representing SD values were added to each distance interval.

### 2.5 Inter-population genomics

Population genetic summary statistics were calculated in GenoDive to assess levels of genetic diversity. These analyses were conducted with full length STACKS loci (i.e. SNPs + monomorphic loci) to calculate heterozygosity independently of global sample size biases (Schmidt et al. 2021). Subsequent analyses were calculated with only the first SNP per locus to avoid linkage disequilibrium among SNPs. First SNPs were directly exported from STACKS or genotype likelihoods were exported from ANGSD to vcftools, filtered there and re-imported into ANGSD. To assess the partitioning of genetic variation between islands, populations, and individuals, a hierarchical analysis of molecular variance (AMOVA; (Excoffier et al. 1992)) was used in GenoDive with an infinite allele model and 999 permutations to assess significance.

To assess levels of population differentiation, pairwise population genetic differences between islands and between populations were calculated using FST, GST and DEST, as recommended by (Verity and Nichols 2014). All three calculations were conducted with GenoDive, estimating significance based on 9,999 permutations with subsequent sequential Bonferroni correction to adjust significance for multiple comparisons. Isolation-by-distance among populations was assessed in GenoDive for all three distance measures (FST, GST & DEST) using a Mantel test (Mantel 1967) with 9,999 permutations. In-water geographic distances among populations were estimated on Google Earth and log converted (Rousset 1997).

To determine and visualize the presence of genetic structure between Saipan and Guam, and among Guam populations, covariance matrices were constructed with the ANGSD subprogram, PCAngsd. The R package “vegan” was then used to convert them for principal coordinate analysis (PCoA) with the constrained analysis of principal coordinates function, as in (Barfield et al. 2020).

To further determine patterns of genetic structure, NGSadmix (Skotte et al. 2013) was used to estimate admixture proportions from the likelihood data. The resulting bar charts were plotted in R, following Skotte et al. (Skotte et al. 2013), for genotypic cluster values K=1-6 to determine genome-wide *A.* cf. *pulchra* admixture.

Migration rates among populations were estimated with BA3-SNPs ((Wilson and Rannala 2003, Mussmann et al. 2019). Total and burn-in iterations were tested to ensure their convergence, and set to 4,000,000 MCMC iterations, 1,000,000 burn-in and sampling interval = 100. Mixing parameters (migration rates dM, allele frequencies dA, and inbreeding coefficients dF) were established empirically as well to obtain an acceptance rate between 20 - 60 % as recommended by BA3-SNPs manual, resulting in the following final parameters settings: dM=0.45, dA=0.95, dF=0.1. Finally, a 95% confidence interval (CI) was constructed as instructed in the program manual (mean ± 1.96 * sdev).

### 2.6 Signatures of selection analyses

Two different FST outlier approaches were used to identify differential selection in pairwise population comparisons. First, we ran BayeScan vs 2.1 (Foll and Gaggiotti 2008) with default parameters and only loci with a q-value below 0.05 were considered statistically significant outliers. Second, we used BayPass Version 2.4 (Gautier 2015), with an ANGSD VCF output that was converted to BayPass format using the reshaper_baypass script by Yann Dorant (gitlab.com/YDorant/Toolbox). BayPass was run once to obtain the covariance matrix between populations (mat_omega), which was then used to control for population structure in a set of 5 independent MCMC runs with different seeds. The median value for XtX over all five runs was then used for each SNP. Additionally, we simulated a neutral distribution of 1000 loci using the simulate.baypass function in the BayPass R script baypass_utils.R and generated 5 independent runs with the same approach as outlined above to obtain their distribution and define the threshold to identify a locus as outlier.

2.7 *Symbiodiniaceae* clade type determination

To infer the presence of different *Symbiodinaceae* genera from holobiont RAD data, we used a method developed by (Barfield et al. 2018). Quality filtered and trimmed ddRAD reads were competitively mapped to transcriptomes of *Symbiodinium, Durusdinium, Cladocopium,* and *Breviolum* with Bowtie2 v2.3.5 with default settings excluding soft matches to determine the predominant symbiont genus in each sample. Transcriptomes for *Symbiodinium* and *Breviolum* were acquired from (Bayer et al. 2012), and transcriptomes for *Cladocopium* and *Durusdinium* were from (Ladner et al. 2012). Resulting SAM files were used to calculate relative proportions of reads with highly unique matches, determined by a mapping quality of 40 or higher to each *Symbiodiniaceae* transcriptome, using the custom perl script zooxType.pl (https://github.com/z0on/).

To verify this ddRAD-based symbiont genus-typing approach, we conducted a ITS metabarcoding approach for a subset of samples. Briefly, we amplified ITS2 following Baumann et al. (Baumann et al. 2018) using the primers SYM_VAR_5.8S2/SYM_VAR_REV (Hume et al. 2018). Amplifications were sent to Azenta Life Sciences for sequencing 2x300 bp paired-end reads on a Illumina MiSeq platform. Raw sequence data are available on the NCBI Sequence Read Archive under the BioProject accession (tbd). The dada2 pipeline (Callahan et al. 2016) in R was used with a reference database that included ITS2 references form *Symbiodinium, Durusdinium, Cladocopium,* and *Breviolum*.

## 3 RESULTS

A total of 267 ddRAD libraries for 249 unique *Acropora* specimens and 18 technical replicates were sequenced to produce more than 200 million raw reads overall. Among these, 233 samples were identified as *A.* cf. *pulchra* in phylogenomic analyses (Fig. S1) and, after quality filtering and removing samples with less than 5,000 high-quality raw reads, 188 *A.* cf. *pulchra* (incl. 18 technical replicates) were used to identify clones.

### 3.1 Clonality, Diversity and Spatial Genetic Structure

For this basic dataset, including potential clones and technical replicates, probabilistic genotype likelihoods generated with ANGSD resulted in 16,780 SNPs, genotyped in at least 50% of samples. Hierarchical clustering analysis based on identity-by-state (IBS) distances in ANGSD detected two clearly different types of IBS-relationships among samples (Fig. S2):

1. Small IBS distances of 0.1-0.2 (average, median and peak = ∼0.15), were found among technical replicates and many intra-population pairs that we consider clone mates. As expected, clone mates had slightly higher genetic distances than technical replicates (Avg. 0.153 vs. 0.147) since somatic mutations accumulate after fragmentation, which may have occurred many years ago.
2. In contrast, IBS-distances between 0.2-0.4 (average, median and peak = ∼0.3) represent relationships among genotypes resulting from sexual reproduction and were found within and among populations. This approach identified 74 unique genotypes, 36 of which had multiple ramets (2 - 19 ramets per genet; Avg. 3.7; Fig. S2, Table 1), so less than half of all samples (44%) constituted unique individual genotypes generated via sexual reproduction.

**TABLE 1:**
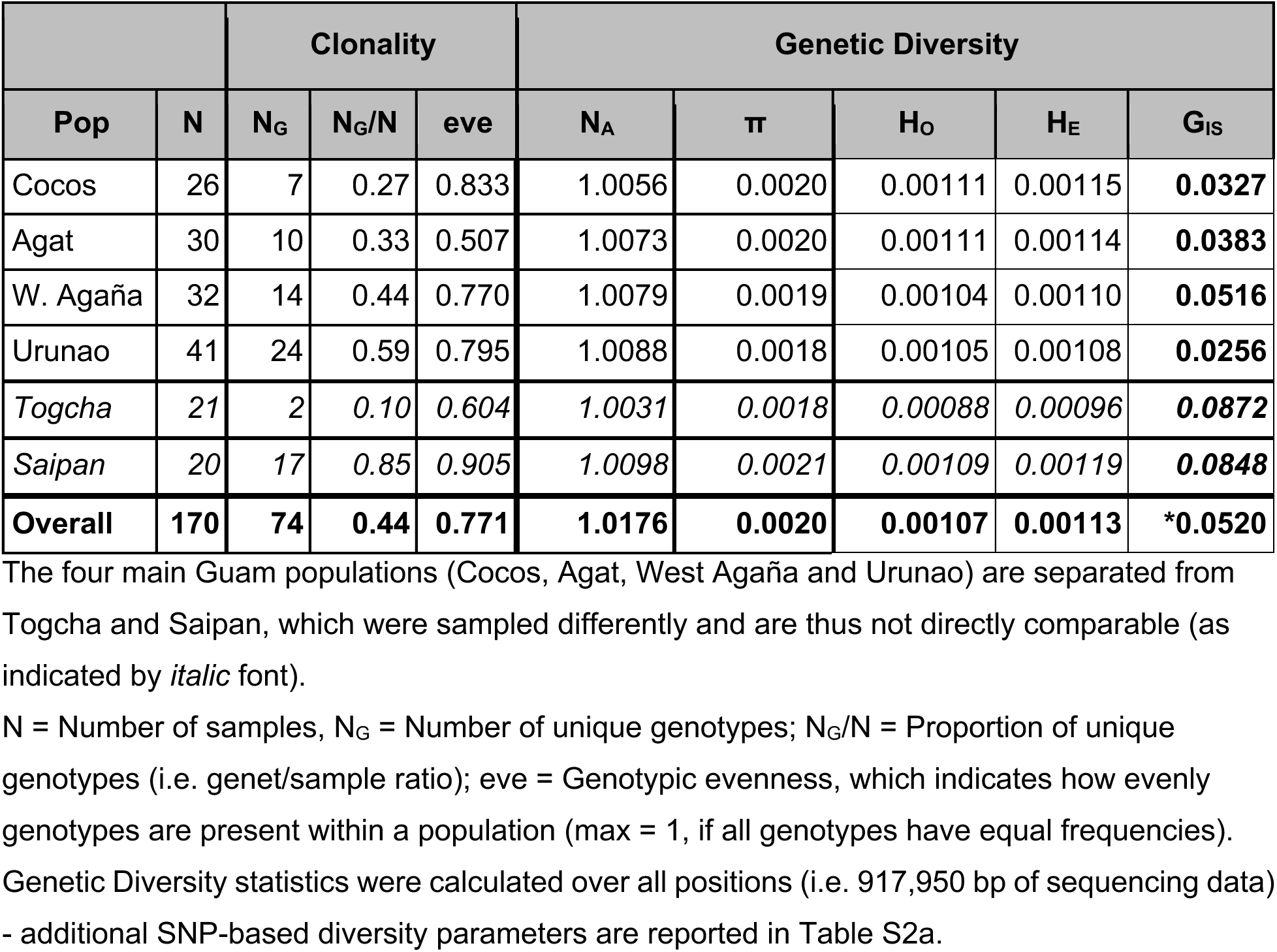
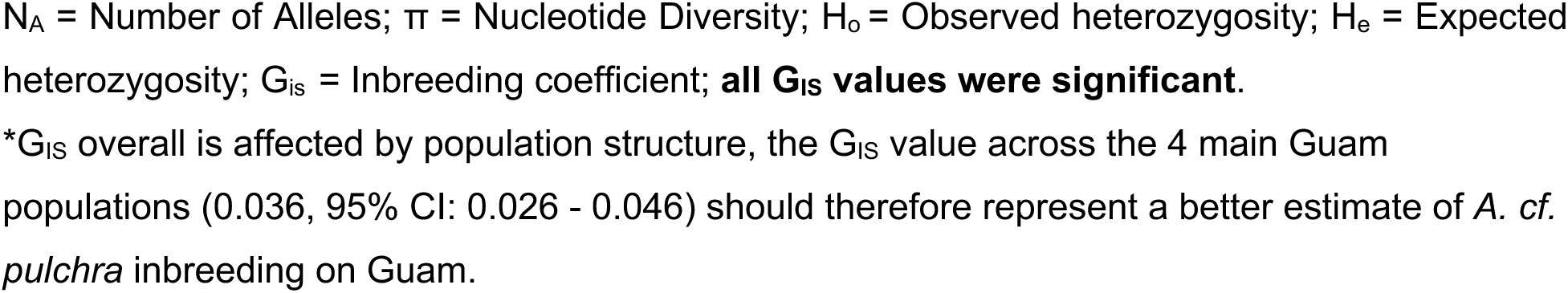
Clonality and Genetic Diversity statistics.

|  |  | Clonality |  |  | Genetic Diversity |  |  |  |  |
| --- | --- | --- | --- | --- | --- | --- | --- | --- | --- |
| Pop | N | $N_G$ | $N_G/N$ | eve | $N_A$ | $\pi$ | $H_O$ | $H_E$ | $G_{IS}$ |
| Cocos | 26 | 7 | 0.27 | 0.833 | 1.0056 | 0.0020 | 0.00111 | 0.00115 | **0.0327** |
| Agat | 30 | 10 | 0.33 | 0.507 | 1.0073 | 0.0020 | 0.00111 | 0.00114 | **0.0383** |
| W. Agaña | 32 | 14 | 0.44 | 0.770 | 1.0079 | 0.0019 | 0.00104 | 0.00110 | **0.0516** |
| Urunao | 41 | 24 | 0.59 | 0.795 | 1.0088 | 0.0018 | 0.00105 | 0.00108 | **0.0256** |
| *Togcha* | *21* | *2* | *0.10* | *0.604* | *1.0031* | *0.0018* | *0.00088* | *0.00096* | ***0.0872*** |
| *Saipan* | *20* | *17* | *0.85* | *0.905* | *1.0098* | *0.0021* | *0.00109* | *0.00119* | ***0.0848*** |
| **Overall** | **170** | **74** | **0.44** | **0.771** | **1.0176** | **0.0020** | **0.00107** | **0.00113** | **\*0.0520** |

The four main Guam populations (Cocos, Agat, West Agaña and Urunao) are separated from Togcha and Saipan, which were sampled differently and are thus not directly comparable (as indicated by *italic* font).
$N_A$ = Number of Alleles; $\pi$ = Nucleotide Diversity; $H_o$ = Observed heterozygosity; $H_e$ = Expected heterozygosity; $G_{IS}$ = Inbreeding coefficient; **all $G_{IS}$ values were significant.**
\* $G_{IS}$ overall is affected by population structure, the $G_{IS}$ value across the 4 main Guam populations (0.036, 95% CI: 0.026 - 0.046) should therefore represent a better estimate of *A. cf. pulchra* inbreeding on Guam.

The proportions of unique genotypes (NG/N; Table 1) showed a strong north to south gradient along the west coast of Guam: the highest proportion of unique genotypes (NG/N = 0.59) was found in Urunao and the lowest in Cocos (0.27). The genotype evenness was fairly high, i.e. clones are rather evenly distributed among genotypes within populations, with the exception of Togcha and Agat, which are dominated by one and two clones, respectively.

Saipan had the highest NG/N (0.85) and Togcha had the lowest NG/N overall (0.10) but since these two populations were sampled haphazardly, their genotypic diversity is not directly comparable. Interestingly, the small and remote population of Togcha was sampled thoroughly but only two unrelated genotypes were identified: one in 19 ramets and the other one in only two (Table 1).

After the removal of clones and technical replicates, 74 samples with unique genotypes comprised the final population genetic dataset (n=74). ANGSD generated 19,940 SNPs for these 74 samples and after the removal of all but one SNP per RAD locus and subsequent VCF filtering, 994 independent SNPs remained for population genomic analyses. For a subset of analyses, fixed genotypes were generated using STACKS, which resulted in 11,490 RAD loci and 25,820 SNPs. Subsequent removal of all but one SNP per RAD locus and further vcf filtering resulted in 1,376 independent SNPs.

Population genetic summary statistics show similarly low levels of genetic diversity among islands and populations (Table 1). The number of alleles (NA) closely follows the number of genotypes per population. Nucleotide Diversity (π) and observed and expected heterozygosity (HO & HE) were virtually identical across populations, with slightly more diversity in Southern Guam (i.e. in Cocos and Agat). Slightly higher diversity metrics were found on Saipan, but differences between Guam and Saipan were small. Inbreeding coefficients were all significantly positive, indicating minor heterozygote deficiency, but levels of inbreeding were low overall (∼0.01 - 0.1). In the ramet dataset (includes clones), Spatial Genetic Structure (SGS) was strong and significantly positive (i.e. beyond the 95% confidence intervals) over the first four distance intervals (i.e. up to 40m; FR10 = 0.183, p = <0.005; Fig. S3). Between 40-180m, SGS was still consistently positive but within the 95% confidence interval, i.e. not significant. The genetic patch size, where colonies are on average as related to each other as the population average, was 180m and the Sp statistic for the ramet dataset was 0.049. In contrast, the genet dataset (i.e. excluding clones) showed no significant SGS: average relatedness values varied randomly around 0 and were mostly within its 95% confidence, i.e. non-significant (Fig. S3). Consequently, the Sp statistic was only 0.0008 only.

### 3.2 Population structure and genetic differentiation

All three measures of pairwise population differentiation indicate small but significant genetic differences between the islands of Guam and Saipan (*F*ST= 0.024, p < 0.001; G’ST = 0.022, p < 0.001; Dest = 0.005, p < 0.001). In addition, all population differentiation measures between individual populations were significant as well, except for Cocos vs. Agat and West Agaña vs. Urunao, i.e. among populations in northern and southern Guam (Fig. 2 & Table S3).

**FIGURE 2:**
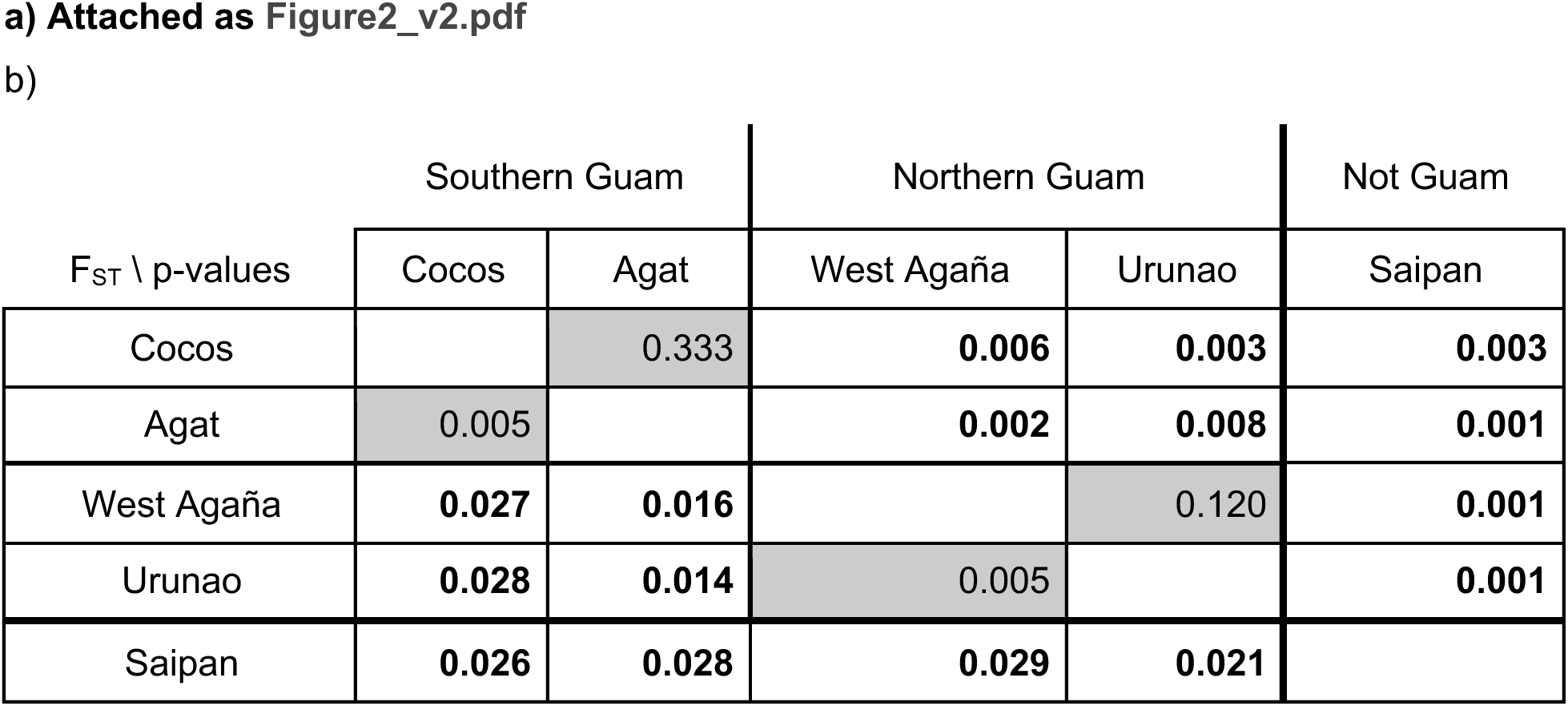
F_ST_ differentiation and Isolation by distance. Pairwise F_ST_ values (below triangle) and associated p-values (above triangle) between populations. All comparisons in bold on white ground have a significant p value after sequential Bonferroni correction (p = 0.05). Togcha was excluded from these analyses due to its low number of unique genotypes (n=2).

Mantel tests confirmed a significant correlation (p < 0.05) of pairwise genetic differentiation (assessed as IbD-transformed FST/(1-FST), sensu (Rousset 1997), as well as untransformed F**ST**, G’**ST** and D**EST**; Table S3) and geographic distances between populations, both regular and log-transformed (e.g. r^2^ = 0.38 - 0.59; p = 0.008 - 0.025; Fig. 2, Table S4). Around Guam, IbD explained an even larger portion of genetic differentiation (r^2^ = 0.77) but was not significant (p = 0.161), presumably due to the low number of comparisons (n = 6) among Guam populations.

In addition to significant overall IbD patterns, there was also a clear pattern of differentiation between Southern (Cocos & Agat) and Northern populations (West Agaña & Urunao), which will subsequently be referred to as the Northern and Southern “metapopulation”. The presence is most evident in the much lower and non-significant pairwise differentiation among populations within metapopulations (Fig. 2) - in contrast to the much higher and significant differentiation among metapopulations. For example, pairwise differentiation between Agat and West Agaña which are located in different metapopulations, is much higher and significant compared to the similarly-distant population pairs within metapopulations (FST = 0.016 over 30 km vs. 0.005 over 16 km between Cocos - Agat, and 0.005 over 22 km between West Agaña - Urunao).

AMOVA analyses confirmed that slightly more genetic variation is partitioned between these two metapopulations (1.6%; p < 0.001) than among the four main Guam populations (1.5%, p < 0.001; Table S5). Hierarchical AMOVA analyses among all populations further confirmed that highly significant proportions of genetic variation are partitioned between islands (1.5%; p < 0.001) and among populations on Guam (1.4%; p < 0.001; Table S5).

The principal coordinate analysis (PCoA; Fig. 3) largely confirms patterns of pairwise differentiation found in the genetic distance metrics. Guam and Saipan separate along the first principal coordinate, but Saipan overlaps significantly with the Northern Guam populations. The two Guam metapopulations are clearly distinct although they overlap substantially and populations within metapopulations overlap almost completely (Fig. 3).

**FIGURE 3:** Principal coordinate analysis (PCoA) **Submitted as Figure3.231209.pdf**

Visual inspections of admixture plots, conducted with NGSAdmix for K=2-6 for all populations and only Guam populations, indicated no clear pattern with increasing K, so only K=2 and K= 3 are included here (Fig. S4). Both admixture plots emphasize the main difference between Guam and Saipan. For example, with K=2, all but one Saipan sample were predominantly affiliated with the “red” cluster (in K=2) and 12 out of Saipan 17 samples showed 100% genetic affiliation with that cluster. In contrast, most Guam samples were dominated by the “green” cluster (i.e. >50%) and 13 out of 57 Guam samples had 100% affiliation with the green cluster. In addition, both plots indicate more admixture from Saipan in Northern Guam populations (Urunao & West Agana), compared to Southern Guam populations (Cocos and Agat). For example, with K=3, 75% of Urunao and 43% of West Agana samples had observable proportions of admixture from the blue Saipan cluster, but only 20% and 14% of Agat and Cocos samples did (Fig. S4).

### 3.3 Relatedness and migration

Relatedness provides a direct snapshot of dispersal and connectivity within and between populations. Since some individual pairwise comparisons did not share many loci, only pairwise comparisons including at least 10% of all RAD loci (100 out of 994) were considered valid. Out of these 2589 pairwise comparisons (Fig. 4), 194 showed elevated levels of relatedness (7.5%), including 157 third-degree relatives (e.g. “cousins”), 35 second-degree relatives (e.g. “half siblings”) and 2 pairs of first-degree relatives (e.g. “full siblings”, one in West Agana and the other split between Cocos and Agat). On average, 11.3% of all intra-population pairs were related, compared to 7.7% among populations and 5.1% among islands. Most intra-population pairs were found in Saipan (25 out of 136 comparisons, 18%) but the highest proportion of relatives was found in Cocos (5/21, 24%), i.e. within the two big lagoon populations.

**FIGURE 4:** Relatedness within and among populations **Submitted as Figure4.240916.pdf**

Inter-island comparisons revealed the highest proportion of relatives for Saipan genotypes were found in the two Northern Guam populations (5.7% with Urunao and 6.4% with West Agana) and Togcha (5.9%). On Guam, inter-population relative pairs were more common within metapopulations than between (10.4% and 7.5%, respectively), especially in the South of Guam (19.1%). The two genotypes in Togcha had relatives in Cocos, West Agana, Urunao and Saipan as well, tentatively confirming its connection with other Guam populations.

Direct estimates of migration rates among the four main Guam populations indicate that populations rely predominantly on self-seeding, with on average 75% of recruits originating from the same population and over 80% in Agat. Inter-population migration rates were unevenly distributed among populations (Fig. 5). Agat was identified as the main source population for inter-population migrants, contributing between 15 - 20% of recruits to other populations. In contrast, migration rates out of Cocos and West Agaña were low (∼2%) with 95% confidence intervals overlapping with zero, indicating an absence of recent migration. These two populations thus may act as larval sink populations.

**FIGURE 5:**
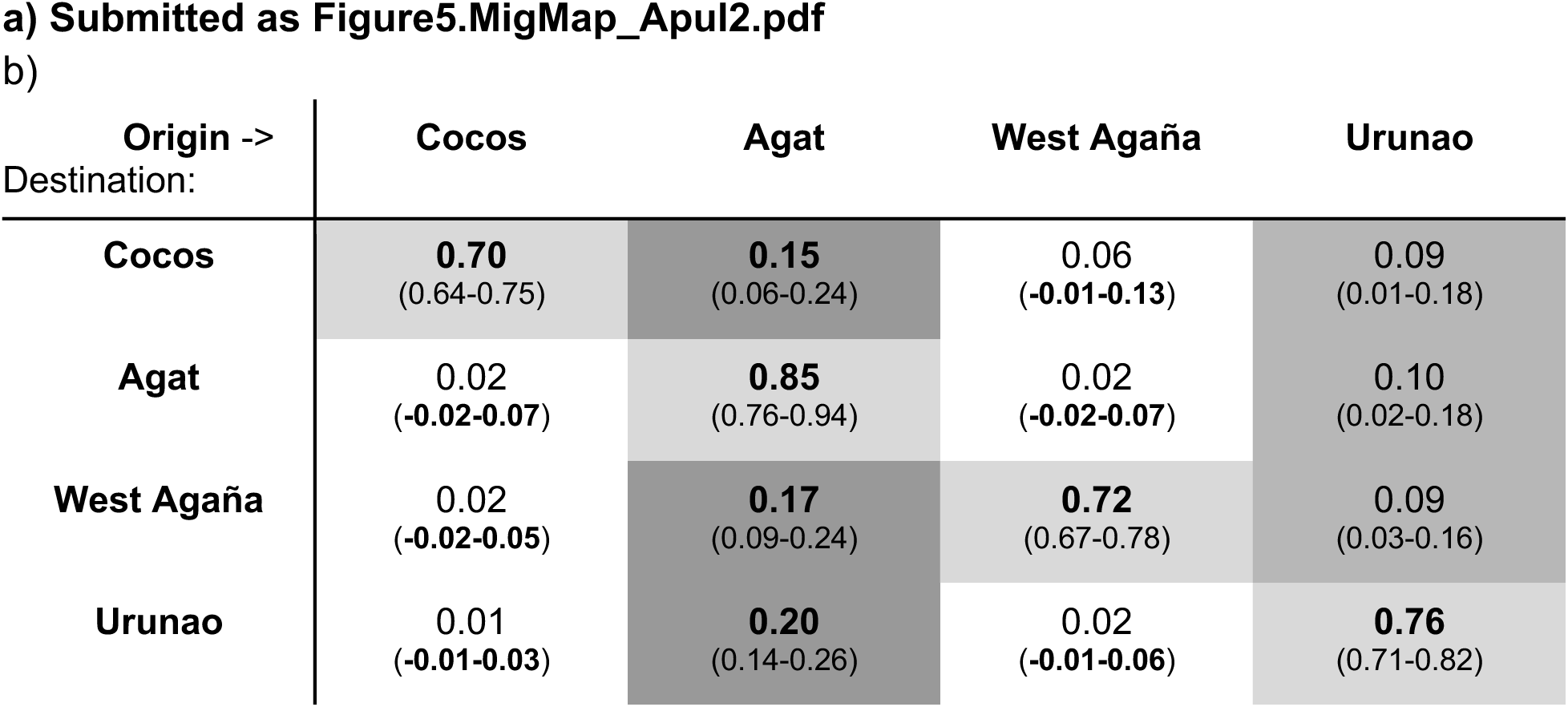
Migration among Guam populations Proportion of individuals in each population that originated in the population itself and in other populations, as calculated with BA3-SNPs.Rows: Assessed population; Columns: Population of origin. Values in brackets indicate 95% confidence interval.

### 3.4 Loci under selection

BayPass and BayeScan were used to detect outlier loci in pairwise population comparisons. BayPass detected a total of 84 different RAD loci under selection, mostly between islands (n = 37, Table 2). Among metapopulations, most putative loci under selection were detected between Northern Guam vs. Saipan (24) and vs. Southern Guam (19), compared to Southern Guam vs. Saipan (12), potentially indicating similar selection regimes in Saipan and Southern Guam.

**TABLE 2:** Number of putative loci under selection between populations.

| Meta-population: |  | South |  | North |  |  | Saipan |  |  |
| --- | --- | --- | --- | --- | --- | --- | --- | --- | --- |
|  | Population: | Cocos | Agat | W. Agaña | Urunao |  |  |  |  |
| South | Cocos |  | 0 | 7 | 2 | 24 | 5 | 15 | 44 |
|  | Agat |  |  | 3 | 9 |  | 3 |  |  |
| North | W. Agaña |  |  |  | 1 |  | 12 | 29 |  |
|  | Urunao |  |  |  |  |  | 10 |  |  |

Number of outlier loci between islands, metapopulations and populations as detected by BayPass. Color scale corresponds to low (white) vs. high (dark grey) values.

Among pairwise population comparisons, 34 putative loci under selection were identified. Among islands, most loci under selection were again detected between Saipan and populations in Northern Guam (12 and 10 loci) compared to Saipan vs. Southern Guam populations (5 and 3).

On Guam, most loci under selection were found in comparison across metapopulations (n = 2-9) compared to within (n = 0-1). Eight loci were found in more than one pairwise comparison and two of them among the same pair of populations, between WAG-Saipan and WAG-Cocos, again indicating potentially similar selection regimes in Saipan and Cocos, the two major lagoons. BayeScan did not detect any significant FST outliers, which is not unexpected since this approach has higher thresholds to indicate significant selection (e.g. (Lotterhos and Whitlock 2015).

### 3.5 Algal symbiont characterization

Photosymbiont communities of *A*. cf. *pulchra* in the Southern Marianas were *surprisingly* diverse and contained a total of 9 different Symbiodinacea genera. Comparisons of ITS-metabarcoding and ddRAD symbiont genotyping showed remarkably consistent results at the genus level: in all 20 samples tested with both approaches the dominant symbiont genus was identified as *Cladocopium*, and most samples had an overall very similar symbiont community composition (Fig. S5). The presence of low frequency background genera, including *Effrenium*, *Freudenthalidium*, *Gerakladium*, (formerly clades E, F, G) and clades H and I (all under 0.02%) were not consistently recognized by either method and individual genera were absent in 7 ddRAD and 6 ITS barcoding results. Therefore, we focus here on the dominant symbiont genera (= >80% of symbiont reads) in the more comprehensive ddRAD dataset.

In total, more than 20,000 ddRAD reads aligned to the four symbiont genomes from 224 different *A.* cf. *pulchra* dataset (90.7 reads/sample on average) and 165 samples had at least 5 reads aligned to symbiont transcriptomes (Fig. 6). All samples were clearly dominated (i.e. >80%) by either *Cladocopium* (C; n = 128, 78%) or *Durusdinium* (D; n = 37, 22%). According to our ITS2 profiling, 89% of *Cladocopium* reads belong to C40 and another 8% could not be assigned to a *Cladocopium* species. For *Durusdinium*, 99% of all reads belonged to D1.

**FIGURE 6:**
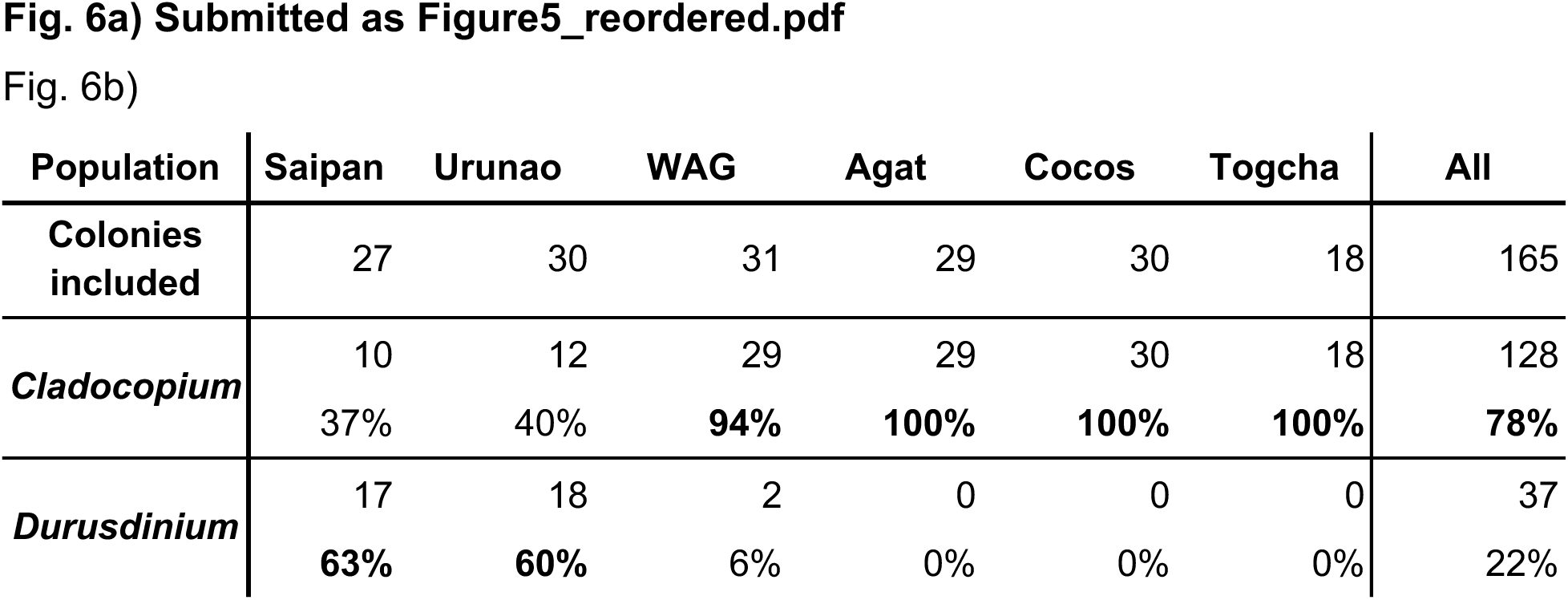
Dominant Photosymbionts The relative proportions of ddRAD reads producing highly unique matches to transcriptomes of four different genera of algal symbionts, Symbiodinium, Breviolum, *Cladocopium*, and *Durusdinium* (formerly Clades A-D, respectively). The distribution of colonies dominated by either *Cladocopium* or *Durusdinium* was significantly uneven among populations (p < 0.0001).

A significantly uneven distribution of dominant symbiont genera was detected among islands and among populations on Guam. On Saipan, most colonies predominantly hosted *Durusdinium* (17/27), while on Guam, most colonies predominantly hosted *Cladocopium* (118/138). Interestingly, 90% of *Durusdinium*-dominated colonies on Guam were found in Urunao, where 18 out of 30 colonies were dominated by D. The two other D-dominated Guam colonies were found in nearby West Agaña, i.e. D-dominated colonies were only found in the two “Northern” populations on Guam (Fig. 6).

Comparisons of symbiont profiles among clonemates (ramets) show that the vast majority of clonemates (101/102 ramets in 22/23 genets) hosted the same dominant symbiont genus. In fact, only one genet in Urunao was dominated by different genera: URU09 had 80% *Durusdinium* reads and only 20% *Cladocopium,* while 100% of all symbiont reads from its clonemates URU08 and URU11 were identified as *Cladocopium* (Fig. 6). This result was confirmed by ITS2 metabarcoding, with 69% *Durusdinium* vs. 31% *Caldocopium* in URU09 compared to 99.9% *Caldocopium* in URU08.

## 4 DISCUSSION

### 4.1 Populations are dominated by large clonal clusters

The first notable finding of our investigation was the high level of clonality in all *A.* cf*. pulchra* populations on Guam and Saipan. Less than half of our samples (74 out of 170) represented unique genotypes. The actual proportion of clones is likely even higher, given that most samples were collected at intervals of 10 meters and the one population sampled more intensively, Togcha, had a significantly higher proportion of clones than any other population (Table 1).

Genotypic diversity on Guam (NG/N = 0.27 - 0.59) is at the lower end of what is commonly observed in highly clonal staghorn thickets like *Acropora cervicornis* (NG/N = 0.17 - 0.71; (Drury et al. 2019) and *A. palmata* (NG/N = 0.51; (Baums et al. 2006) but see (Japaud et al. 2015)). Lower levels have occasionally been detected in other coral species but different sampling designs make direct comparisons difficult (e.g. in *Pocillopora damicornis*: (Combosch and Vollmer 2011) vs. (Torda et al. 2013a) vs. (Gorospe and Karl 2013) vs. (Adjeroud et al. 2014)). Here, clone mates were only found within populations (Fig. S2, Table 1), indicating vegetative fragmentation as the predominant and likely sole source of clonality (Tunnicliffe 1981, Highsmith 1982). High levels of clonality indicate limited contributions of sexual reproduction to population maintenance, which is in line with the low fecundity observed in *A.* cf. *pulchra* populations around Guam (Lapacek 2017, Raymundo et al. 2022) and with generally extremely limited coral larvae recruitment in the Mariana Islands (Birkeland et al. 1981, Neudecker 1981, Minton et al. 2007). Low reproductive output may be a consequence of environmental stress and degradation since reproductive capacity is one of the first processes to be compromised when corals are stressed (Ward 1995, Ward et al. 2000, Baird and Marshall 2002). In addition, recent mortality events severely reduced the number of *A.* cf. *pulchra* colonies (Raymundo et al. 2017, 2019) and thus genotypes, i.e. potential mates, which may further interfere with sexual reproduction (Ortiz et al. 2018).

The spatial distribution of *A. cf. pulchra* clones on Guam is characterized by tight clusters of clones and some exceptionally distant clone mates. The significant SGS pattern in the ramet dataset (i.e. including clones), but not in the genet dataset (i.e. excluding clones), indicates that the presence of clonal clusters significantly increases the average relatedness up to 40 m around colonies. The *A. cf. pulchra* ramet Sp statistic of 0.049 (a metric to quantify and compare SGS) is similar to estimates for *P. damicornis* populations in the Tropical Eastern Pacific (TEP; 0.055, (Combosch and Vollmer 2011) but higher than in a more clonal Hawaiian population (0.005, (Gorospe et al. 2015). This indicates that *A.* cf. *pulchra* clones are strongly clustered on Guam, en par or more so than in other clonal coral populations. Despite these generally tight clonal clusters, several clonemates were separated by over 100m and one clonal pair in Agat was 200 m apart. Ramets of other staghorn species, e.g. *A. palmata* and *A. cervicornis*, are usually only up to 25 m and 75 m apart, respectively (Baums et al. 2006, Japaud et al. 2015, Drury et al. 2019). Larger distances between clonemates have been observed in *Pocillopora* corals but these have been attributed to well-dispersed asexual larvae (which have never been observed in *Acropora* corals) rather than fragmentation (Souter et al. 2009, Torda et al. 2013b, Adjeroud et al. 2014, Gélin et al. 2017). The abundance and significant spatial extent of *A.* cf. *pulchra* clones indicates a long and successful history of clonal lineages, especially in Agat, where clones were most distant and clonal evenness was particularly low (Table 1). Nonetheless, the low levels of genotypic diversity are concerning for the adaptive capacity of these threatened populations in the face of environmental change (Jump et al. 2009, Pauls et al. 2012).

In contrast to the significant ramet SGS, the genet SGS and Sp statistic for *A.* cf. *pulchra* were extremely low (0.0008), indicating that sexual recruitment is basically random within populations and sexually-derived, related colonies are not clustered within populations - as expected for broadcast spawners (Stoddart 1988, Miller and Ayre 2008) and lower then for example in most terrestrial plants (Sp = 0.0003 - 0.04)(Vekemans and Hardy 2004, Dering et al. 2015). Nonetheless, several closely related colonies were found within populations, particularly in Cocos and Saipan (Fig. 4), which is a hallmark of sweepstake reproductive successes as a consequence of broadcast spawning (Barfield et al. 2022) and likely indicates retention of larvae during development within these two big lagoons (Fig. 1). Development times for *A. pulchra* on the GBR are estimated to be 7-10 days (Baird 2001), while larvae of the closely related *A. millepora* (Torrado et al, 2024) may settle within 3 days (Connolly and Baird 2010). Given the typically calm conditions during *A. cf. pulchra* spawning in May (pers. observ.), it seems plausible that larvae are retained within large lagoons for the entire 3-10 day developmental period. This is supported by our findings of widespread self-seeding (Fig. 5) and likely contributes to the observed population structure.

### 4.2 Past and present signs of vulnerability

Overall, our results uncovered numerous signs that the Guam *A. cf. pulchra* populations are vulnerable to further degradation. Not unexpected but most concerning is the extremely low genetic diversity in all populations: As discussed above, the genotypic diversity of *A. cf. pulchra* thickets on Guam is low, which leads to low resilience in the face of abiotic and biotic disturbances (Reusch et al. 2005), especially disease epidemics (Vollmer and Kline 2008). In addition, levels of nucleotide diversity (π) are among the lowest recorded in any coral species so far and are ∼10- 100 times below levels observed in ecologically and phylogenetically similar *A. cervicornis* in Florida reef tract (Drury et al. 2017) or *A. tenuis* on the GBR (Matias et al. 2022). The nucleotide diversity of *A. hyacinthus* in nearby Yap is most comparable, but still 2-3 times higher (Barfield et al. 2022). Nucleotide diversity is often used as a measure of evolutionary potential (O’Grady et al. 2004, Kardos et al. 2021), which seems severely compromised for *A. cf. pulchra* on Guam. Likewise, observed and expected heterozygosities are also well below commonly observed levels (e.g. (Sole-Cava and Thorpe 1991, Hellberg 2006, Hemond and Vollmer 2010, Drury et al. 2017). While this is not entirely unexpected for small populations on remote oceanic archipelagos that suffered significant recent mortality (Raymundo et al. 2019), it is concerning for their persistence and adaptive capacity (sensu (Haig 1998, Reed and Frankham 2003, Oppen and Gates 2006, DiBattista 2008, Shearer et al. 2009). In addition, small and isolated populations are more susceptible to further loss of genetic diversity due to limited gene (in)flow, genetic drift, limited opportunities for sexual reproduction and recurrent bottlenecks (e.g. (Noreen et al. 2009, Robinson et al. 2016). Together with the recent declines in overall abundance and the loss of several local populations, this indicates a need for urgent intervention, e.g. genetic rescue (as discussed below).

### 4.3 Connectivity among staghorn populations is limited

The dominant feature of the population genetic structure among *A. cf. pulchra* populations are the significant patterns of isolation-by-distance (IbD, Fig. 2): depending on the exact parameter of differentiation, between 38-59% of the total genetic variation among populations can be explained as a function of geographic distance between populations. Excluding Saipan, even more genetic differentiation was explained by the geographic distance among the four main Guam populations (77%) but IbD was not significant (p = 0.16), presumably due to the lower number of pairwise comparisons (n = 6). Isolation by distance is a direct result of limited dispersal over the analyzed geographic distances, i.e. the observed IbD patterns suggest that *A.* cf. *pulchra* larvae do not effectively disperse between Guam and Saipan (sensu (Wright 1943, Kimura and Weiss 1964, Slatkin 1993). Although IbD among Guam populations was not significant, the significant population structure indicates that populations are not well connected around Guam either, i.e. over 10s of kilometers. The steeper IbD slope (Fig. 2) further indicates that pairwise differentiation on Guam increases faster over shorter distances, which is most likely due to additional factors limiting connectivity, like environmental heterogeneity, e.g. near-shore hydrodynamic forces (Meirmans 2012). Although significant IbD patterns are frequently observed in corals (but see e.g. (Ayre and Hughes 2004, Magalon et al. 2005, Nakajima et al. 2009, Combosch and Vollmer 2011), most studies tend to find weaker patterns over much larger geographic distances, especially for broadcast spawners like *Acropora* corals. For example, (Davies et al. 2014) found isolation-by-distance patterns for *A. digitifera* and *A. hyacinthus* across the Caroline Islands with 62% and 74% of genetic variation explained by geographic distances over ∼4000 km, respectively (but see (Cros et al. 2016)). Other examples include *A. millepora* along the Great Barrier Reef (IBD = 54% over 1550 km; (Oppen et al. 2011)) or *Porites lobata* among Hawaiian islands (IBD = 37% over 2500 km; (POLATO et al. 2010) *but see (Tisthammer et al. 2020)*). The tight IBD pattern observed here, compared to other studies, suggests that connectivity among *A.* cf. *pulchra* populations is exceptionally robust to offshore current patterns and environmental heterogeneity over moderate spatial distances around Guam and Saipan.

A direct consequence of the overall IbD pattern is the significant inter-island genetic differentiation between *A.* cf. *pulchra* populations on Guam and Saipan (e.g. Fig. 2, 3 & 4). There are three other islands, almost perfectly in line between Guam and Saipan: Tinian and Aguijan, which are ∼5 and ∼30 km south of Saipan, and Rota, which is ∼130 km south of Saipan and ∼90 km north of Guam. The central location of Rota presumably facilitates connectivity among the Southern Mariana islands by providing a vital stepping-stone for gene-flow between Guam and Saipan (Fig. 1). Oceanographic measurements and modeling indicate highly variable currents between Guam and Saipan, and predicted most larvae are likely swept westward due to the dominant North Equatorial Current or may be retained locally by leeward eddies (Suntsov and Domokos 2013, Kendall and Poti 2015). In addition, these models identified a clear breakpoint in connectivity between Guam and Rota for larvae with a < 20 day pelagic larval duration (PLD)(Kendall and Poti 2015). The maximum competency period of *A.* cf. *pulchra* is 14 days with settlement often occurring 10 days after fertilization (Baird 2001, Baird et al. 2009), which is in line with only occasional larval exchange between Guam and Rota (Kendall and Poti 2015). Although the role of Rota as a stepping-stone could not be verified here directly, it is likely vital in connecting Guam to any other Mariana Island. This hypothesis is e.g. tentatively supported here by the fact that Urunao, the northernmost population on Guam and only ∼65 km south of Rota, is most connected to Saipan (e.g. Fig. 2, 3, 4 & S4).

On Guam, 8 out of 10 pairwise FST comparisons were significant, and a significant proportion of the genetic diversity is partitioned by population (Fig. 2). Although geographic distances are the strongest predictor for genetic differentiation among populations (Fig. 2), there is also a significant differentiation between Northern (Urunao & West Agaña) and Southern (Agat & Cocos) Guam populations, as e.g. clearly indicated by pairwise population differentiation (Fig. 2 & Table S3), the PCoA (Fig. 3), the elevated number of relative pairs among vs. within metapopulations (Fig. 4), and was confirmed to be substantial and significant in multiple AMOVA analyses (Table S5b). The two metapopulations diverge between West Agaña and Agat, where the coast is dominated by two prominent peninsulas that enclose Apra Harbor (Fig. 1), which breaks the otherwise continuous fringing and lagoonal back reefs along the west coast of Guam and seems to constitute a barrier to dispersal. The differentiation between northern and southern sites may be partially driven by leeward coastal eddies that form off the northern and southern tip of Guam due to the westwards flowing ECC (Wolanski et al. 2003, Storlazzi et al. 2009, Suntsov and Domokos 2013, Kendall and Poti 2015). These eddies are likely important for larval retention on Guam (Kendall and Poti 2014) and form an onshore current that diverges into a south- and a north-bound near-shore current near the center of the west coast of Guam, i.e. where the two metapopulations diverge (Wolanski et al. 2003). Interestingly, BayPass selection analyses identified several putative loci under selection between Northern and Southern populations (5.25 loci on average) but only a single locus between populations within each metapopulation (Table 2). This indicates that the differentiation between Northern and Southern Guam may be enhanced by non-neutral forces, i.e. selection. This hypothesis is supported by the distribution of their photosymbionts: algae that belong to the genus *Durusdinium* are common and frequently dominant in *A.* cf. *pulchra* colonies in the two Northern populations but uncommon and never dominant in colonies in the South (Fig. 6). One potential driver of this differentiation are the significant geologic and hydrologic differences between the physiographic Northern and Southern Guam provinces that are separated by the Pago-Adelup fault, which perfectly aligns with the genetic break between West Agana and Agat: The northern half of the island is an uplifted karst plateau formed on reef-lagoon deposits while the southern half is uplifted volcanic terrain (e.g. Fig. S6, (Taborosi et al. 2004, 2013)), which may lead to environmental differences between metapopulations that drive differential adaptations in lagoon and back reef corals like *A.* cf. *pulchra*.

Second, Agat seems to be an important source population, connecting the northern populations with southernmost Cocos. This hypothesis is based on the migration analysis, which identified Agat as the most important source of larvae (besides self-seeding) for both southern and northern populations and supported by its overlap with other Guam populations in the PCoA (Fig. 3) and the relatedness results, where Agat shares the highest proportion of relatives with other populations (8.1% of all pairwise comparisons). Interestingly, Agat is also the population with the highest heterozygote deficit (besides Togcha), which aligns with the prediction that migration into Agat is low. This is surprising at first since Agat is clearly part of the southern metapopulation - genetically (e.g. Fig. 2, 3 & 4, Tables 1, 2 & S4), geographically (Fig. 1), physiographically (Fig. S6, (Taborosi et al. 2004, 2013) and hydrodynamically (Wolanski et al. 2003). However, increased dispersal from Agat could be explained by the particular and complex near-surface currents and off-shore eddies systems around Guam (Cowen et al. 2000, Wolanski et al. 2003, Kendall and Poti 2014, 2015, Limer et al. 2020, Lindo-Atichati et al. 2020).

### 4.4 Photosymbiont communities

In contrast to most coral host population genetic aspects, the photosymbiont communities of *A*. cf. *pulchra* in the Southern Marianas are surprisingly diverse. Although it is not uncommon for *Acropora* species to host multiple different *Symbiodinaceae* genera and/or species (Oppen et al. 2001, Ulstrup and Oppen 2003, Rouzé et al. 2019), the diversity of Symbiodiniaceae in *A.* cf. *pulchra* on Guam is surprisingly high (e.g. (Rouzé et al. 2019) - especially compared to the exceptionally low host genetic and genotypic diversity. Nonetheless, all colonies were clearly dominated by either *Cladocopium* or *Durusdinium* with a striking north-south gradient of prevailing *Durusdinium*-dominance in colonies from Saipan and Urunao and *Cladocopium*- dominance in all other populations (Fig. 6). Although the two dominant species detected here, C40 and D1, are rather thermo-tolerant (e.g. Jones et al. 2008; Qin et al. 2019), *Durisdinium* is generally considered to be more tolerant to warm water temperatures than *Cladocopium* (Stat et al. 2008, Oliver and Palumbi 2009, 2011, Ladner et al. 2012, Keshavmurthy et al. 2014, Silverstein et al. 2017, Barfield et al. 2018). The dominance of *Cladocopium* in *A.* cf. *pulchra* in Southern Guam (Fig. 6) may thus partially explain the higher bleaching incidents there (Raymundo et al. 2017), compared to the *Durusdinium*-dominated *A.* cf. *pulchra* in Saipan (Lyza Johnston, pers. comm.). Population genetic datasets are particularly suitable to test the relationship between host genotype and photosymbiont communities by comparing clone mates, which enables assessing the stability and/or flexibility of symbiont association over decadal time scales (Baums et al. 2014, Manzello et al. 2019). Here, comparisons indicate that *A.* cf. *pulchra* symbiont associations on Guam are remarkably stable over time and across intra-populational environmental gradients (since almost all clonemates hosted the same dominant symbiont type). The presence of different dominant photosymbionts among one set of clonemates does, however, indicate some flexibility. This could be due to different dominant photosymbionts in different parts of the same colony (e.g. (Rowan et al. 1997) before fragmentation. Alternatively, one of the ramets may have shuffled its dominant symbiont genus post-fragmentation (Buddemeier and Fautin 1993, Baker 2003, Jones et al. 2008, Zhu et al. 2022), e.g. following recent bleaching events (Raymundo et al. 2017). This flexibility has major implications for coral restoration since photosymbionts are essential for the survival of the coral holobiont (e.g. (Falkowski et al. 1984, Muscatine et al. 1984, Baker et al. 2013, Matthews et al. 2017) and the composition of photosymbiont communities can have a major impact on the survival of corals in stressful conditions (Baker et al. 2004, Rowan 2004, Berkelmans and Oppen 2006, Thornhill et al. 2014, Parkinson et al. 2015, Levin et al. 2016, Qin et al. 2019). Previous studies have shown that the presence of stress-tolerant symbiont populations may improve adaptive capabilities and could fuel adaptation through natural or assisted transfer of symbiont among conspecifics (Dixon et al. 2015, Anthony et al. 2017, Morikawa and Palumbi 2019, Schoepf et al. 2019).

### 4.5 Implications for management and restoration

#### Protection & Management

The conservation of existing diversity should always take precedence over restoration and this study provides important guidelines for informed protection and management of this keystone reef-builder. For example, the overall significant isolation-by-distance pattern (Fig. 2) indicates that *A. cf. pulchra* is not able to effectively disperse between Guam and Saipan. Since Rota is the only shallow water habitat between Guam and Aguijan, Tinian and Saipan, and the only island within ∼150 km around Guam, it is likely vital for the connectivity and thus the maintenance of genetic diversity in *A. cf. pulchra* among the Southern Mariana Islands. Personal observations in 2022 indicate that the Rota population is small, marginal and highly unstable with significant recent mortality as indicated by extensive stands of dead staghorn skeleton. It should thus become a high priority for monitoring and protection while or even before its significance for inter-island connectivity can be tested explicitly.

On Guam, our results suggest that Agat and Urunao are particularly valuable for local management and protection. Both populations have a slightly elevated genetic diversity, in terms of genotypic (Urunao) and allelic diversity (Agat)(Table 1) and represent both, the Southern and Northern Guam metapopulations, including their unique standing genetic variation and putative metapopulational adaptations (Table 2). In addition, migration and population structure analyses indicate that Agat is the central hub for population connectivity and gene flow among populations and Urunao is vital for the genetic connection between Guam and other Mariana islands (Fig. 4 & 5). Maintaining both genetic diversity and population connectivity, is vital for species conservation and management (Sala et al. 2002, Palumbi 2003, Hellberg 2007, Jones et al. 2009, Christie et al. 2010, Leiva et al. 2022). The Agat population is currently protected within the War in the Pacific National Park but enforcement is limited and significant threats include sedimentation and nutrient inflow from two nearby “rivers” or stormwater drainages and recent bleaching-induced mortality events killed nearly ⅓ of its staghorn population and completely destroyed the *A* cf*. pulchra* population on neighboring Alutom Island (Raymundo et al. 2022). The Urunao population is currently not protected and threatened by the recent development of a massive military installation (“Camp Blaz”) on a nearby karst cliff and recurrent plans for further coastal development. Two nearby populations of staghorn corals recently disappeared completely (Raymundo et al. 2022)., highlighting the need for protection, e.g. by extending the nearby Ritidian Wildlife Refuge, and stopping further development in this remote corner of Guam.

#### Restoration

Restoration is frequently criticized for trying to restore populations before the causes of the decline have been removed (e.g. (Hughes et al. 2023, Streit et al. 2024). While there are certainly situations and restoration projects that deserve criticism, this overly general rejection is both misleading and cynical, since global climate change and thus reef decline, will continue for decades to come, even if CO2 emission would be eliminated tomorrow. Moreover, it also fails to recognize that important lessons have been learned and restoration practices are constantly being optimized (e.g. (Suggett et al. 2023, 2024, Burdett et al. 2024, Edwards et al. 2024). In addition, decimated populations continue to adapt, and may adapt even faster due to the brutal selection regimes during heat-related population bottlenecks (Smith et al. 2013, Precht and Aronson 2016, Eakin et al. 2022, Lachs et al. 2023), which are then selectively included in coral restoration projects (e.g. (Bowden-Kerby 2022). But corals need to be better protected (as discussed above) and restoration needs to be done in accordance with all available information, especially on small and remote oceanic islands like Guam since there is extremely little room for mistakes.

Carefully selecting fragments for propagation, restoration and captive breeding programs is vital since the stock defines the genetic make-up of restored populations (e.g. (Reynolds et al. 2012, Koch 2021, Nef et al. 2021). The observed high clonality and their significant spatial extent is reassuring for coral restoration on Guam, which so far relies heavily on asexual fragmentation (Raymundo et al. 2022). To recreate current levels of genotypic diversity in restored populations, our results suggest that at least half of all colonies in restored populations may be clones, i.e. derived via asexual fragmentation from existing colonies. However, the partial correlation of elevated clonality with higher bleaching induced mortality (discussed before) suggests that there are risks associated with highly clonal populations and a healthy mix of unrelated genotypes will be an important goal for restoration. To source unique genotypes for restoration efforts, colonies should generally be sampled at least 30 m to ideally 50 m apart, as indicated by the spatial genetic structure results (Fig. S3). Nonetheless, clones should be spread throughout restored populations, to increase chances of outbreeding and thus successful sexual reproduction and reduce local inbreeding due to non-random mating within populations (i.e. FIS, Table 1 and below). The observed gradient in clonality suggests that populations in the northern part of the island are more valuable as sources of fragments for asexual propagation (Table 1). The *A.* cf. *pulchra* population in Urunao is particularly attractive due to its elevated genotypic diversity (Table 1), high proportion of colonies with thermotolerant *Durusdinium* photosymbionts (Fig. 6), and likely adaptations to the particularly harsh environmental conditions in shallow back reefs (Table 2). To increase genetic diversity and avoid outbreeding depression, stock colonies should also be sourced elsewhere, explicitly including the genetically distinct but genotypically less diverse Southern populations (Table 1, Fig. 3 & S4).

Increasing the very low genetic diversity of *A. cf. pulchra* in the Southern Mariana Islands is vital for the long-term survival of the species in this region. Genetic diversity can be increased by boosting the remaining diversity and/or by bringing in new genetic diversity from elsewhere (i.e. genetic rescue, (Whiteley 2015,Bell 2019)). Boosting the local genetic diversity is generally preferable to preserve local adaptations (e.g. (Tallmon 2004) and the remaining *A.* cf. *pulchra* genotypes on Guam seem exceptionally temperature-tolerant (Combosch et al, in prep; Reuter et al., in prep), which may have been shaped by past (Cybulski et al. 2024) and recent (Raymundo et al. 2019) mortality events, exerting strong selection pressures (Smith et al. 2013, Precht and Aronson 2016, Eakin et al. 2022, Lachs et al. 2023). Well-adapted, stress-tolerant and genetically diverse populations could thus potentially be generated if the regional effective population size of *A. cf. pulchra* could be increased to optimize the preservation and use of the remaining local diversity (Libro and Vollmer 2016, Muller et al. 2018, Baums et al. 2019). Increasing effective population sizes can be achieved by decreasing inbreeding and increasing the number of genotypes that participate in sexual reproduction. Here, inbreeding was detected in all populations (F/GIS, Table 1 & S2) as well as between (i.e. as FST) most populations, metapopulations and islands. Inbreeding indicates non-random mating over meter scales and limited dispersal, which could potentially be alleviated if genotypes would be more mixed within and among populations, metapopulations and islands, e.g. by introducing other genotypes into large clonal clusters (Fig. S3), via coral restoration. Since we found indications for differential adaptations among metapopulations and islands (Table 2), genotypes should be monitored carefully for survival and differential performance in different locations. Selection analyses further indicate that signatures of divergent selection are lower between Saipan and Southern Guam, which suggests translocation trials between Cocos and Saipan lagoon would be a promising starting point. Translocations of genotypes across metapopulations and islands would further be beneficial to introduce *Durusdinium* symbionts to Southern Guam (which may lead to occasional symbiont switches as inferred for URU09, as discussed above, Fig. 6). All of these suggestions can be achieved with current asexual propagation approaches, which also provide a safeguard against losing propagated genotypes to genetic drift. Ultimately, however, coral restoration via sexual reproduction would be the most direct, efficient and fastest way to improve the genetic diversity and survival of *A. cf. pulchra* in the Mariana Islands.

To a priori assess the suitability of different reef sites for restoration, numerous factors need to be taken into account that are beyond the scope of this study (see e.g. (Vaughan 2021), for a recent review). Here, we focus on (1) the importance of the site for the connectivity and genetic diversity of *A. cf. pulchra*, (2) the need of the potentially remaining population for anthropogenic intervention and restoration and (3) their suitability for restoration. Based on these considerations, our results and *A. cf. pulchra* survey data over the last 10 years (Raymundo et al. 2019), we identified three local priority areas for coral restoration:

1. The *A. cf. pulchra* population in Agat is particularly important as a connectivity hub (Fig. 5; as discussed above) but experienced significant recent mortality (>50%) and the nearby Alutom staghorn population already disappeared completely (Raymundo et al. 2019, 2022), indicating a clear need for intervention in this area. The deeper, sandy backreef and the protected location inside the National Park make this a very suitable site for successful restoration.
2. The lagoons between West Agana and Agat (Fig. 1) connect the Northern and Southern Guam metapopulations and several small *A. cf. pulchra* populations in this area (not included in this study) have experienced significant recent declines (Raymundo et al. 2022). For example, small *A. cf. pulchra* population in Luminao reef, right next to the opening of Apra harbor where the two metapopulations presumably separate, recently declined severely and would benefit from targeted restoration (Raymundo et al. 2022). Guam’s oldest coral nursery and several active outplanting sites near the Piti bomb holes highlight the area’s appeal and suitability for restoration.
3. To support and connect the only staghorn population on Guam’s east coast, the area between Cocos lagoon and Togcha is significant for Guam’s reefs (Fig. 1). Several *A. cf. pulchra* populations recently disappeared here (Raymundo et al. 2022) and the extremely low, remaining genetic diversity in Togcha (Table 1) clearly indicates that targeted restoration is urgently needed to support this isolated outpost. The shallow back reefs and harsh conditions on Guam’s east coast make restoration more challenging here and may be most suitable for seeding sexually generated spat and recruits in the future. Meanwhile, asexual restoration could focus on Achang, the nearest *A. cf. pulchra* populations further south, which also experienced significant recent declines (>90%; (Raymundo et al. 2022), but is located in a more protected lagoon and Marine Preserve.

## Conclusion

Here, we conducted a comprehensive population genomic assessment of *A.* cf. *pulchra* in the Southern Mariana Islands (Micronesia), to guide management and restoration and as a blueprint for conservation and restoration genomic studies elsewhere. We specifically assessed the following vital aspects and recommend a subset of them for future studies:

a. Clonality and its spatial patterns: Clonality is particularly important for coral restoration programs that rely heavily on asexual propagation, as most coral restoration projects still do (Koch 2021). Here, the highly clonal nature of *A. cf. pulchra* populations on Guam testifies to the general suitability of this approach for restoration but observations of elevated mortality in more clonal populations hint at the limitations of this approach. Moreover, the abundance and distribution of clonality within and among populations provided valuable suggestions, like where and how fragments for propagation should be harvested and replanted to efficiently maximize genotypic diversity. A comprehensive assessment of clonality is therefore highly recommended, esp. if vegetative fragmentation is part of the restoration strategy (Vaughan 2021).
b. Genetic diversity: The genetic diversity of restoration target species is the primary determinant of its recovery and evolutionary potential (O’Grady et al. 2004, Hughes et al. 2008, Drury and Lirman 2017, Kardos et al. 2021) and adaptive capacity (e.g. (Haig 1998, Reed and Frankham 2003, Oppen and Gates 2006, DiBattista 2008, Jump et al. 2009, Shearer et al. 2009, Bailey et al. 2009, Sgro et al. 2011). Here, the extremely low genetic diversity of *A. cf. pulchra* in the Southern Marianas indicates that the restoration and maintenance of genetic diversity should be a major target for restoration to be successful long-term. Measuring genetic diversity to preserve and enhance it are therefore key concerns for any coral restoration program (Reynolds et al. 2012, Koch 2021, Nef et al. 2021).
c. Population structure and distribution of related individuals among populations: Assessing connectivity among natural populations is important to identify source populations and locations for management, conservation and restoration. Here, the significant population genetic structure indicated potential benefits of translocations for restoration to maximize genetic diversity and potentially stimulate sexual reproduction. Knowledge about population structure is particularly valuable to prioritize specific sites but otherwise, may be of limited importance from a purely restoration genetics perspective ().
d. Signatures of selection: Understanding the patterns and the extent of local adaptations is particularly useful to plan and assess the prospects of translocations among populations, e.g. to counter the effects of limited genetic diversity. Here, we found indications for limited, localized adaptations, in particular between Guam metapopulations, which warrants closer monitoring of translocated colonies. Their inconsistent detectability, using different approaches, indicates that the strength of local adaptations is likely limited, i.e. does not preclude translocations. Signatures of selection are therefore useful but unlikely to be the most important parameter to assess in future studies, especially if tracking and monitoring of outplants is planned anyways.
e. Dominant photosymbionts: Since the photosymbiotic communities of corals are major determinants of the holobionts’ thermal tolerance, knowledge about their dominant lineages can be useful to restore more thermal tolerance coral populations. Here, the heterogeneity and uneven spatial distribution of dominant photosymbionts offers new opportunities to incorporate symbiont associations in restoration planning and spread more tolerant symbionts. Assessments of the symbiont community are particularly useful if coral species or genera are flexible in their symbiont association.

The results of our study presented here highlight the necessity to conduct thorough genetic analyses to obtain a clear picture of the complex life history of the coral populations to restore and will hopefully serve as a blueprint for similar restoration genomic studies in other important coral species around the world.

## Data Archiving Statement

Data for this study will be available at the National Center for Biotechnology Information (www.ncbi.nlm.nih.gov/).

## Acknowledgement

This research was funded by the National Science Foundation, award No. OIA-1457769 and OIA-1946352. We are thankful to M. Gawel and Guam’s National Park Service for sampling permits and Dr. Lyza Johnston for providing samples from Saipan. The sampling site at Agat is located within the boundaries of the War In The Pacific National Historical Park and was sampled with the appropriate permit (#WAPA-2018-SCI-0010). All other populations were sampled under a scientific collection permit issued by the Guam Department of Agriculture to the Marine Laboratory at the University of Guam.

---

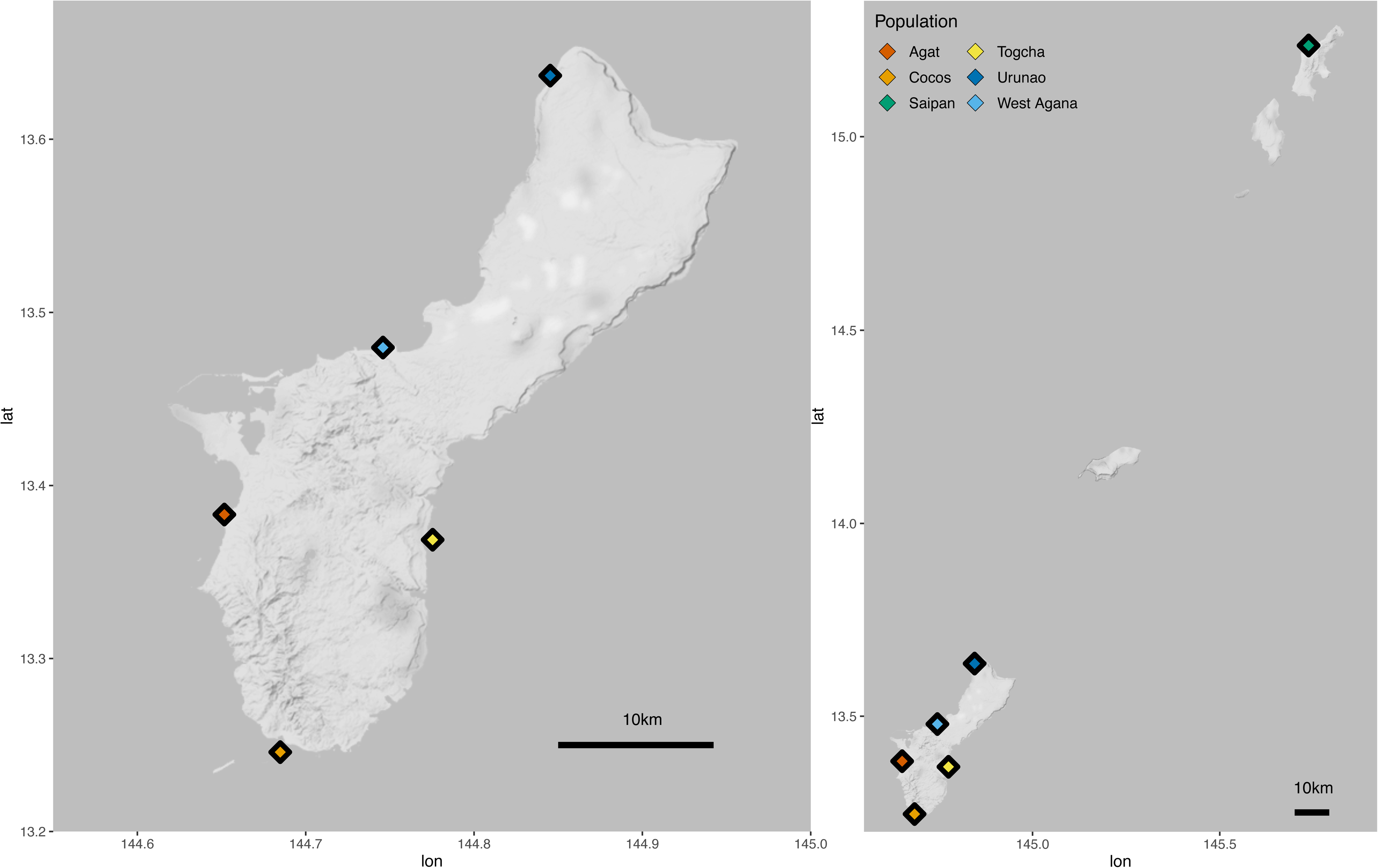

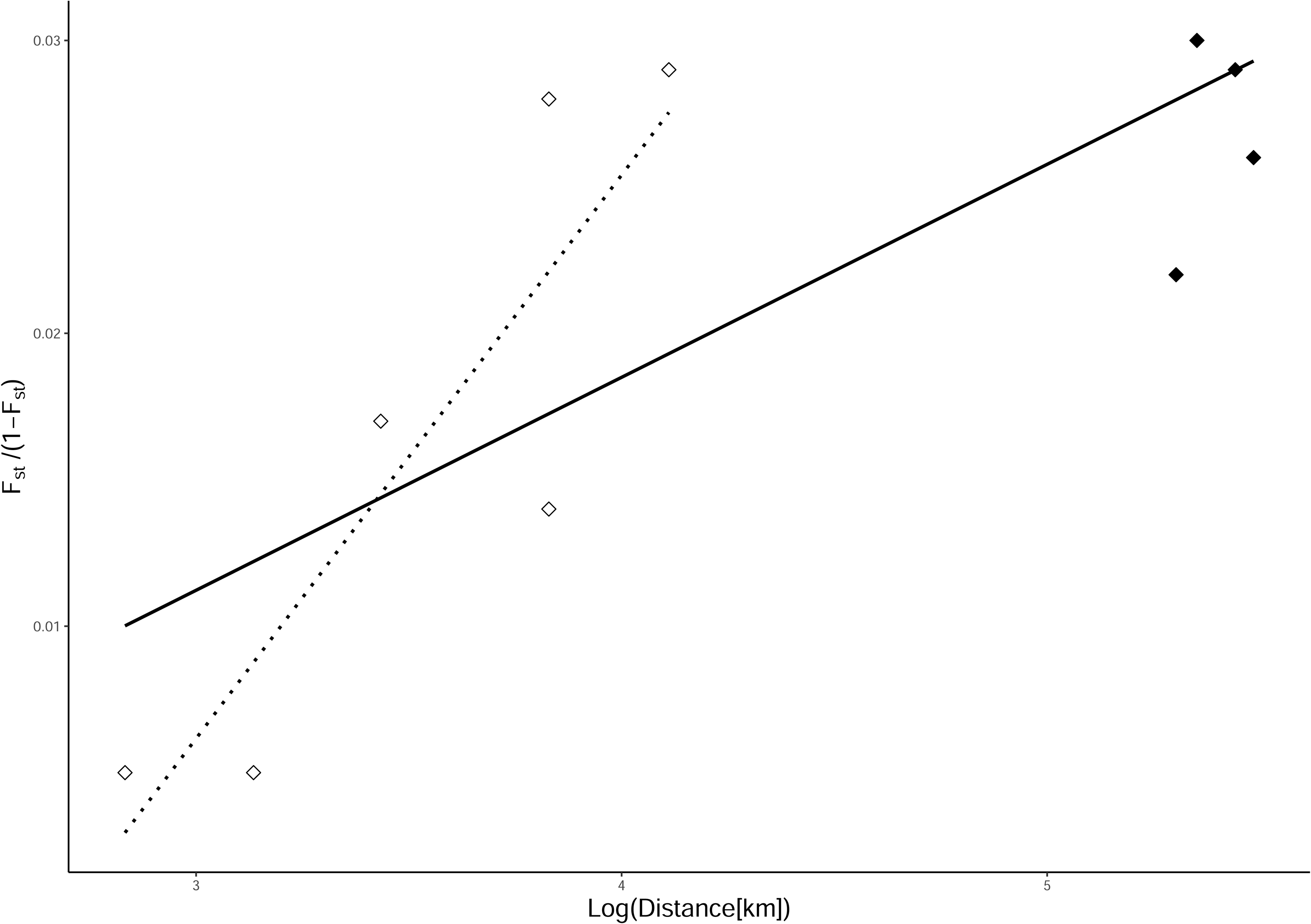

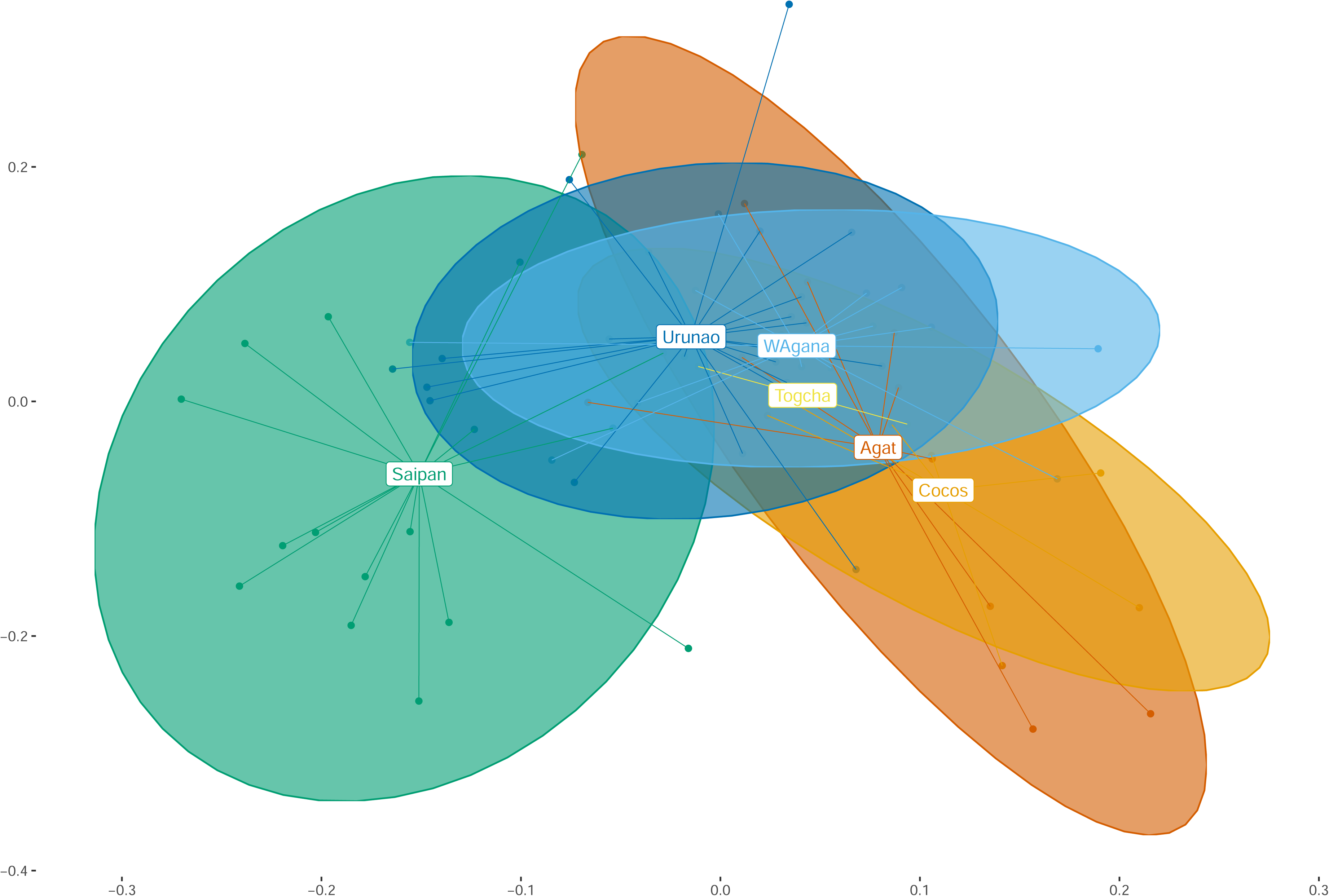

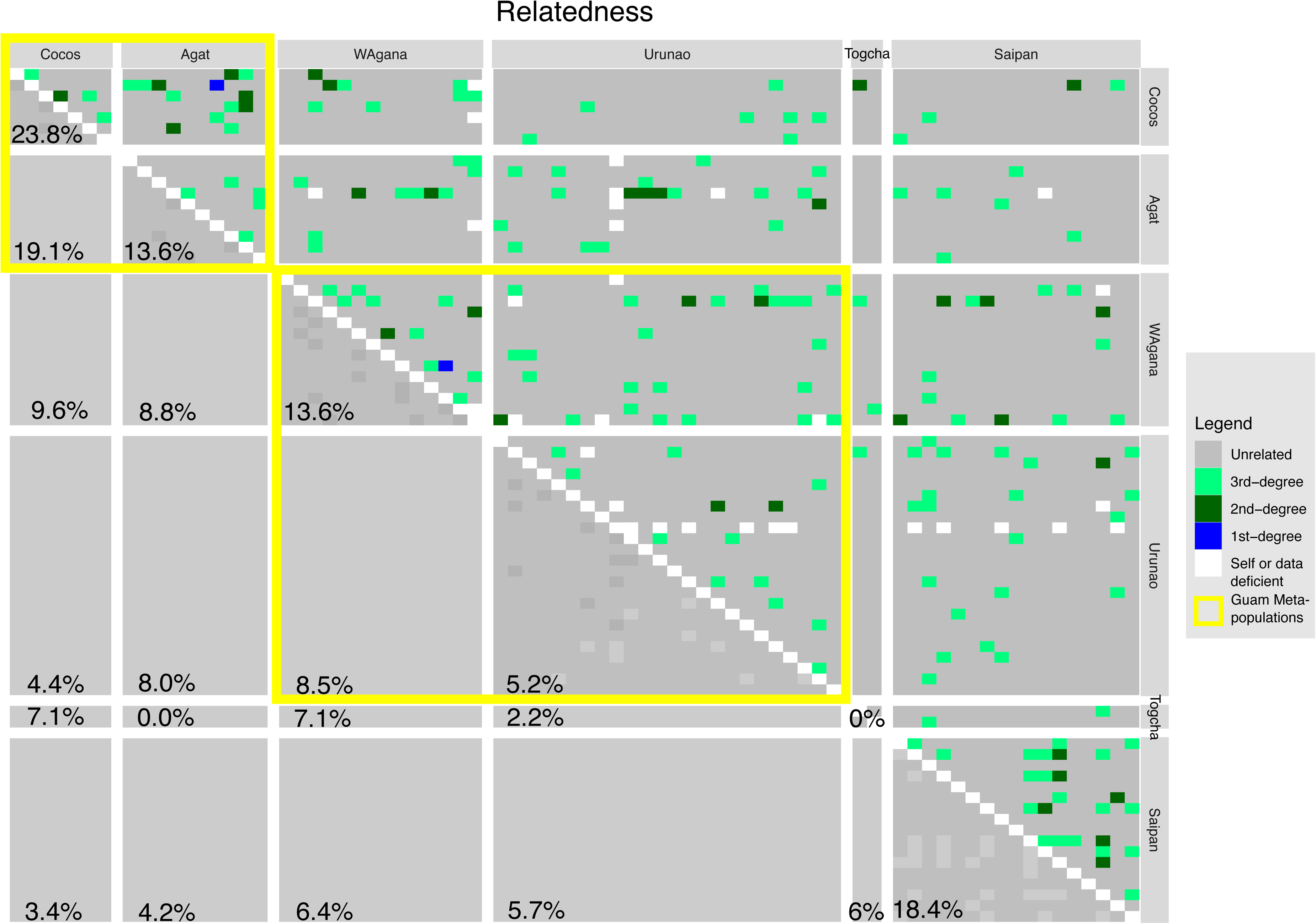

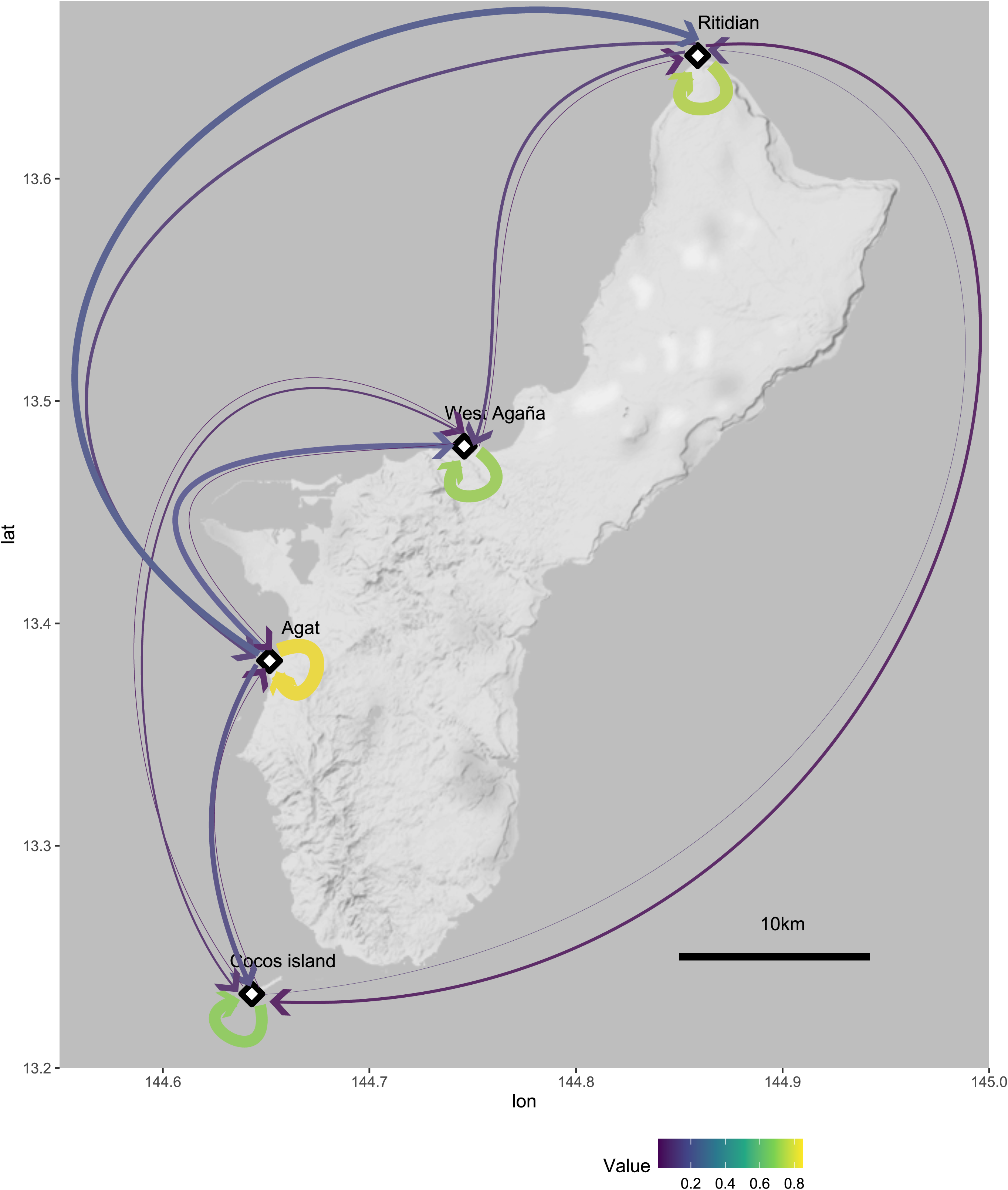

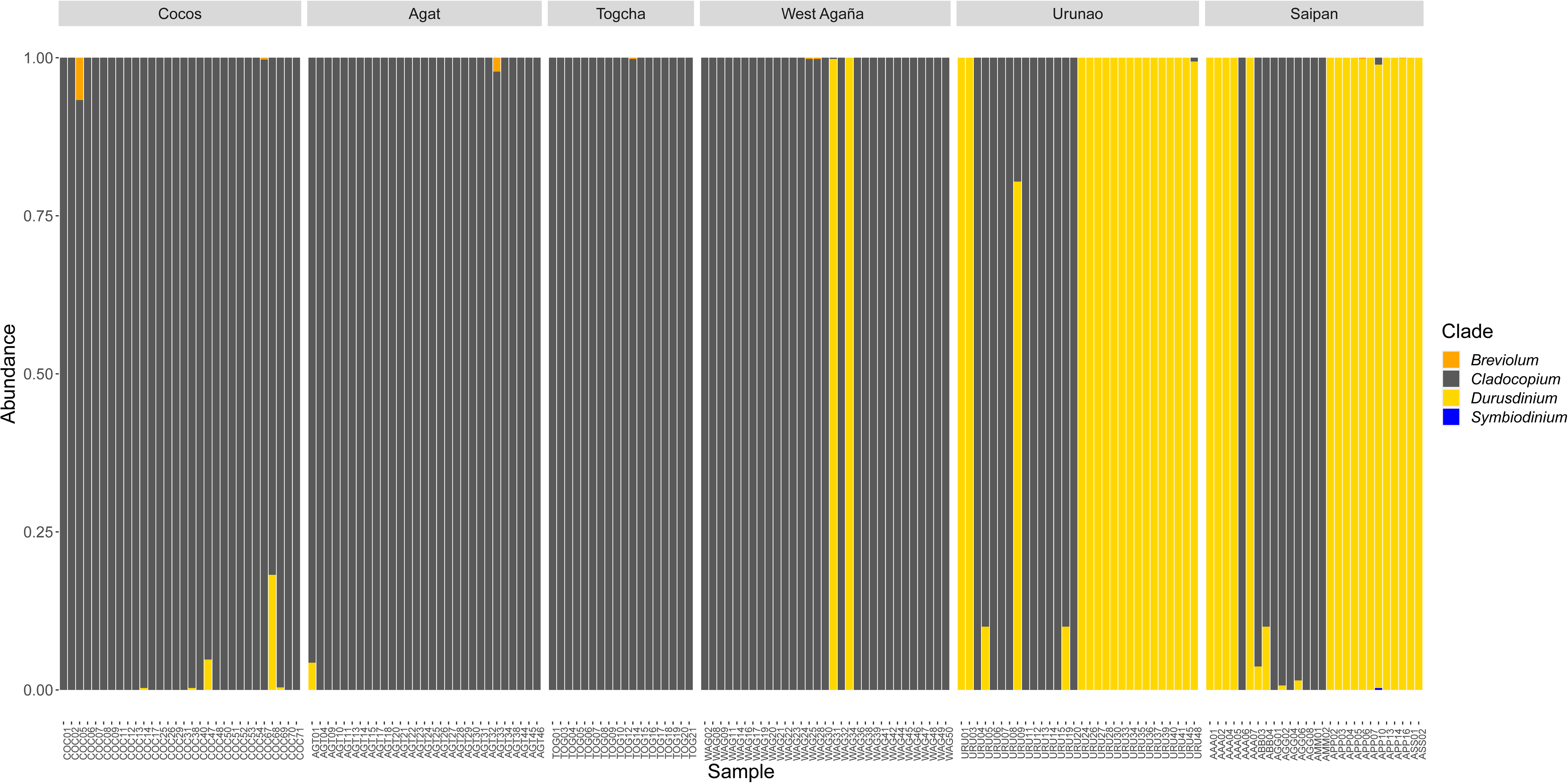

