## Supplementary figures and images for "Population genomics for coral reef restoration - a case study of staghorn corals in Micronesia"

### Fig.S1

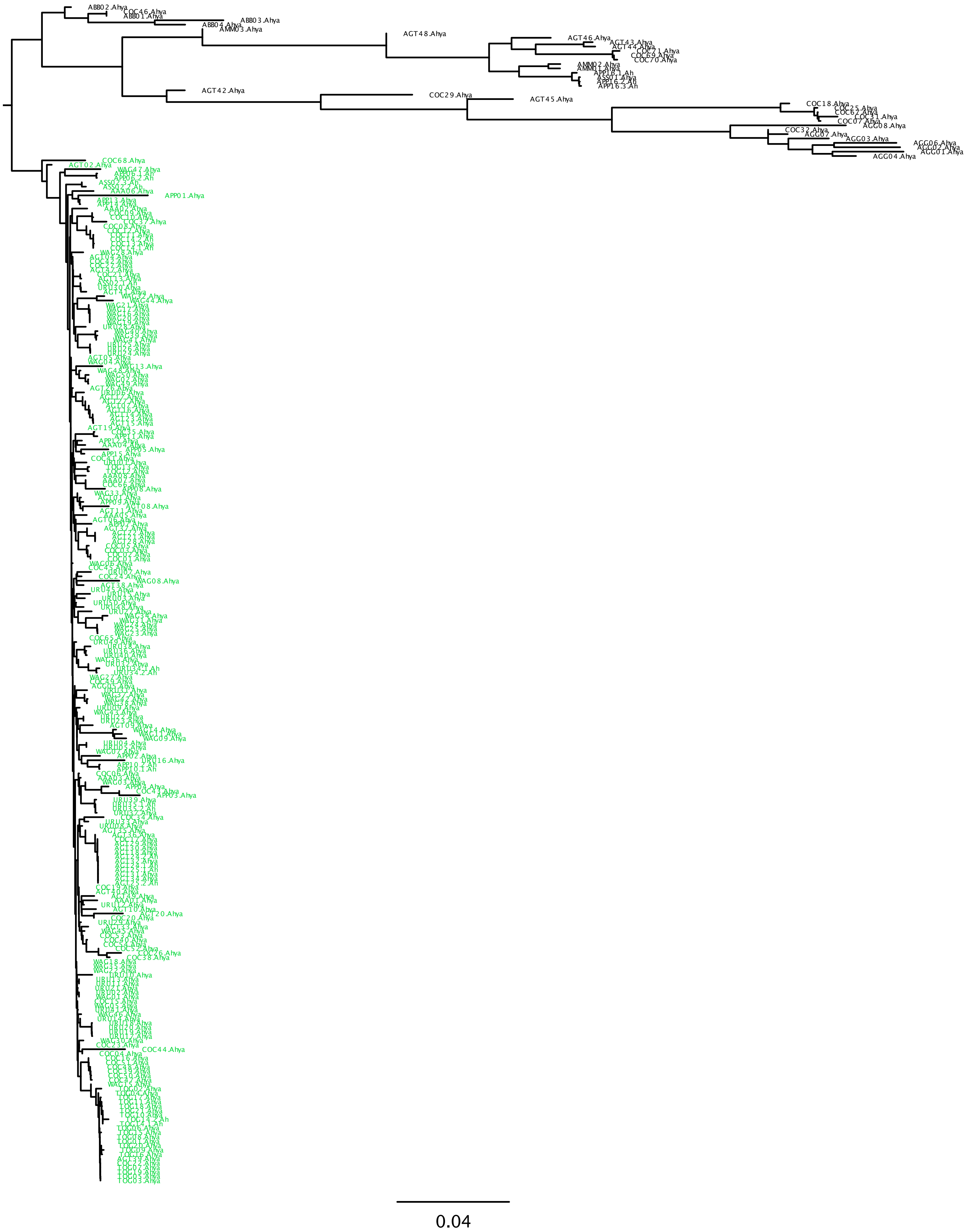

### Fig.S2a

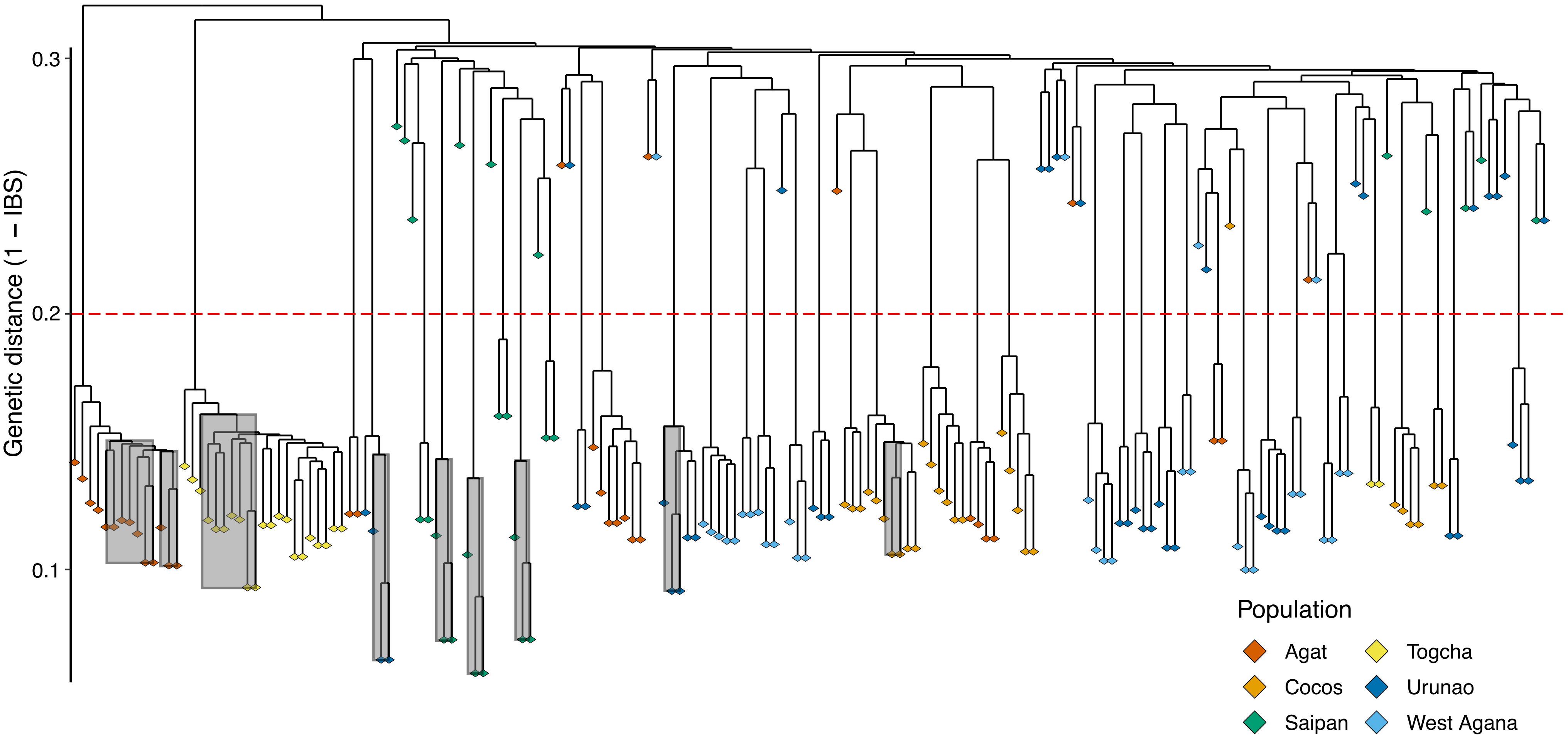

### Fig.S2b

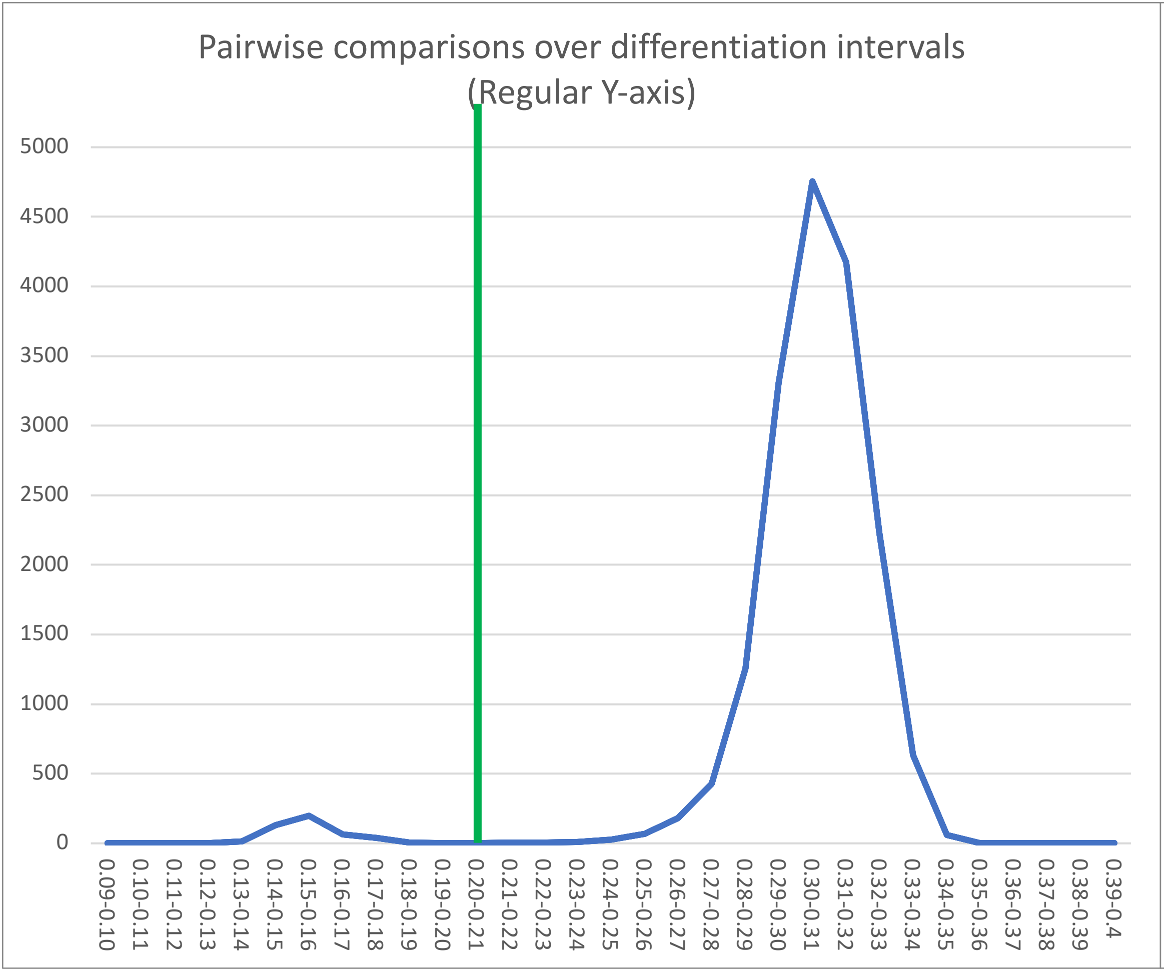

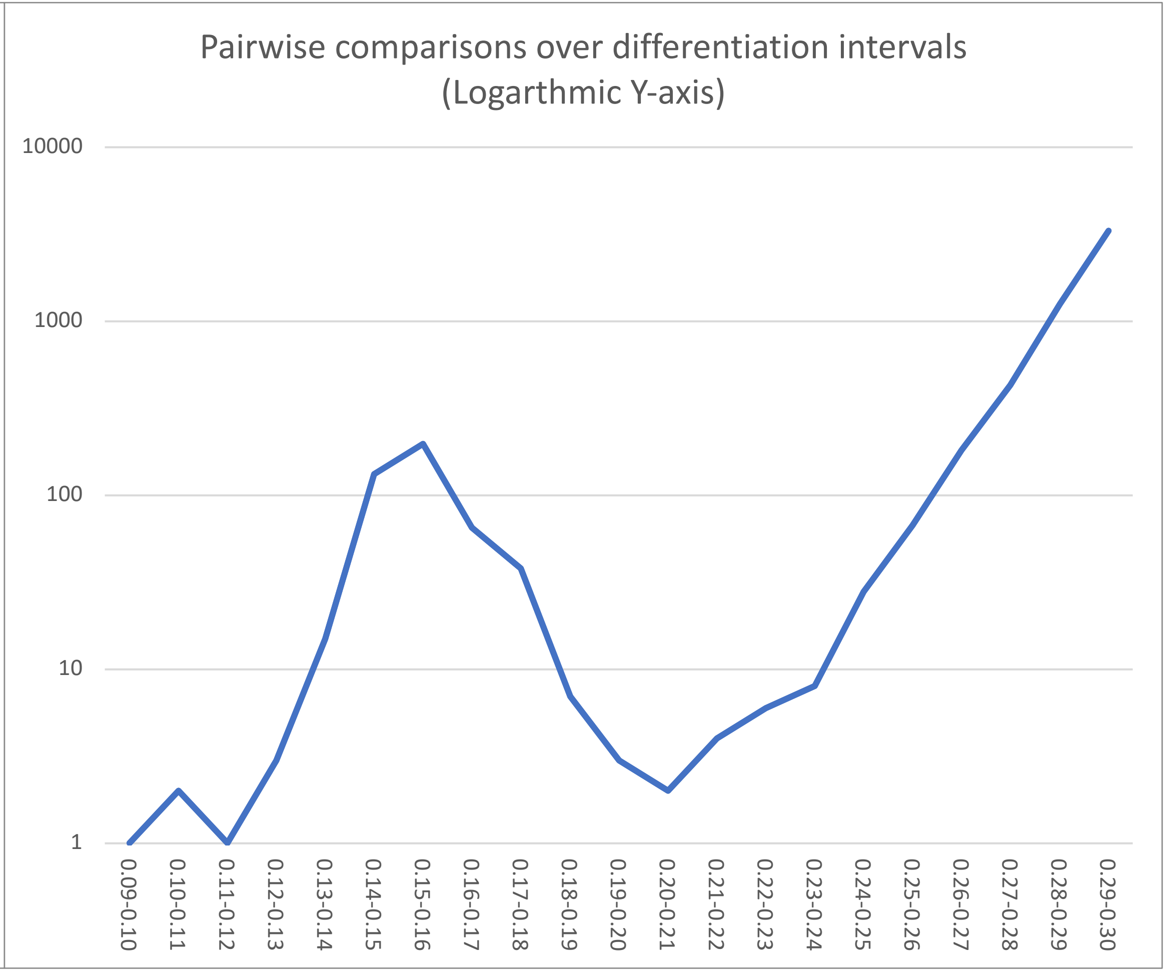

### Fig.S3

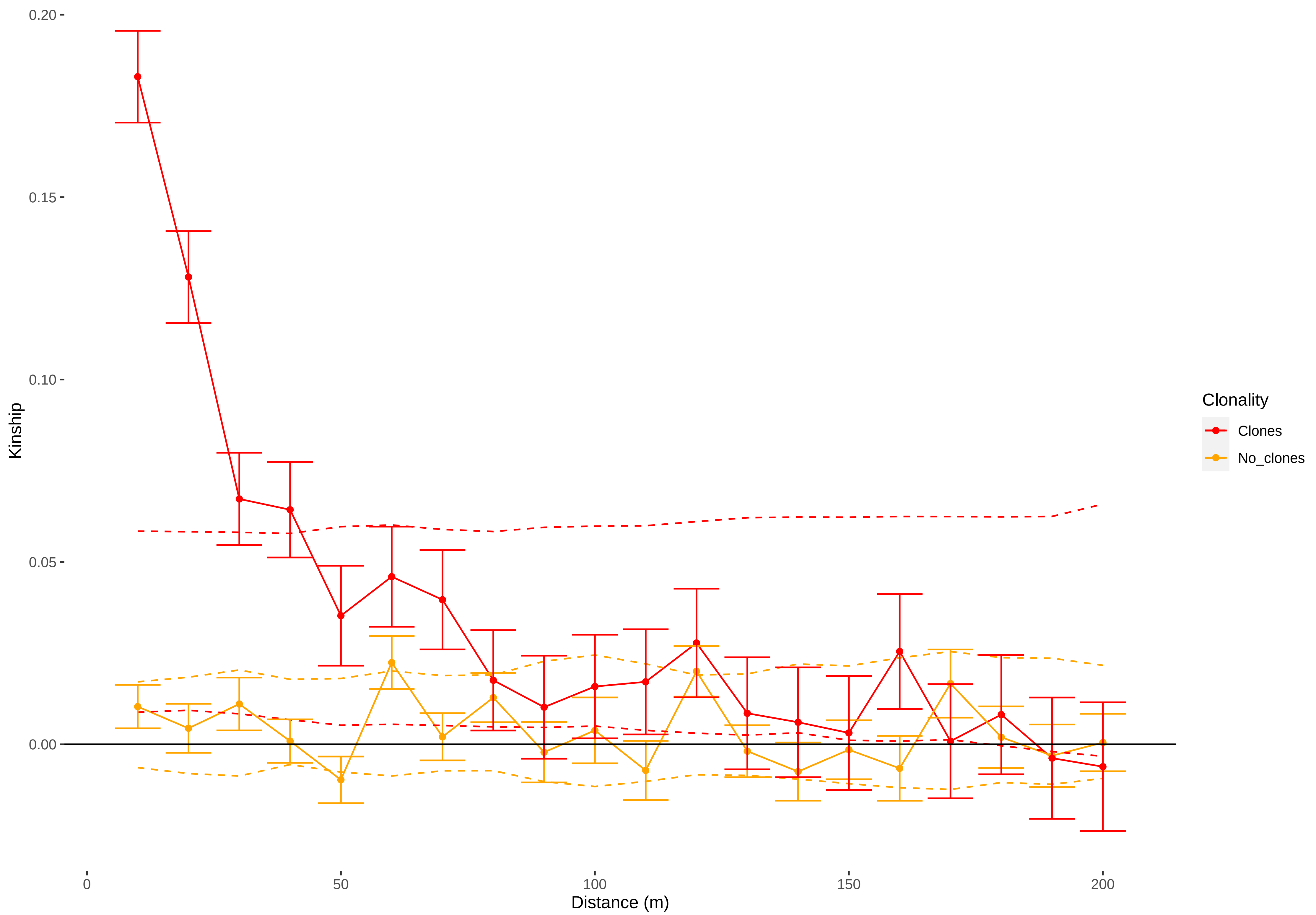

### Fig.S4

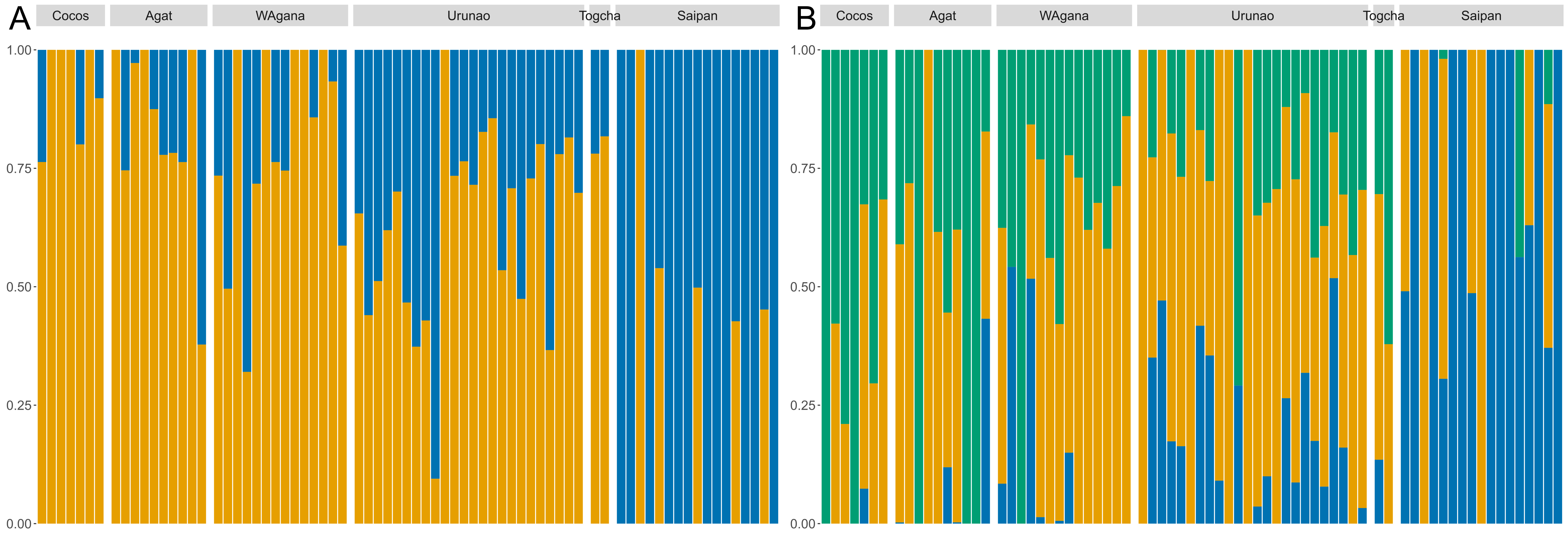

### Fig.S5

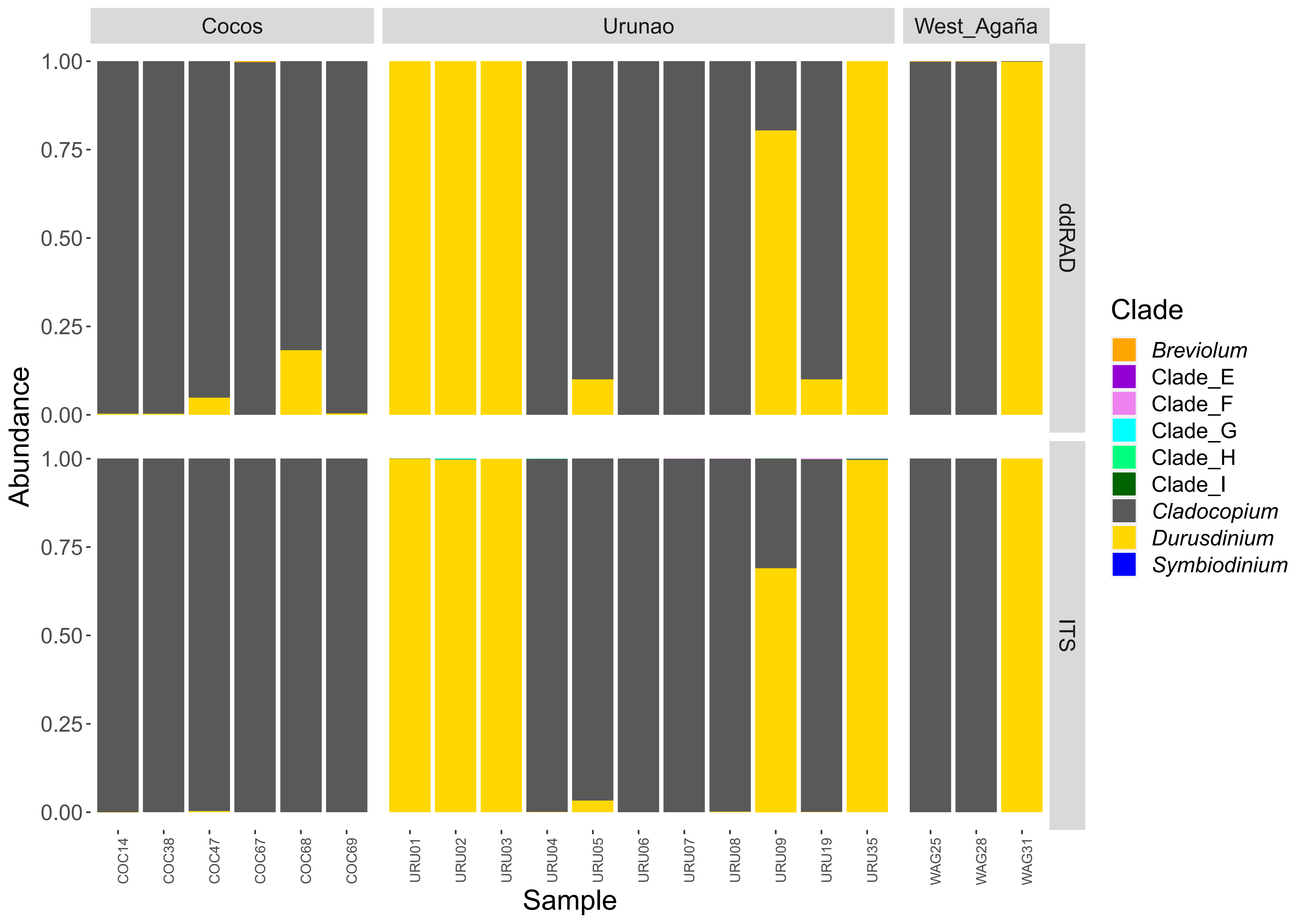
