## Supplementary material for "Population genomics for coral reef restoration - a case study of staghorn corals in Micronesia": Fig.S6

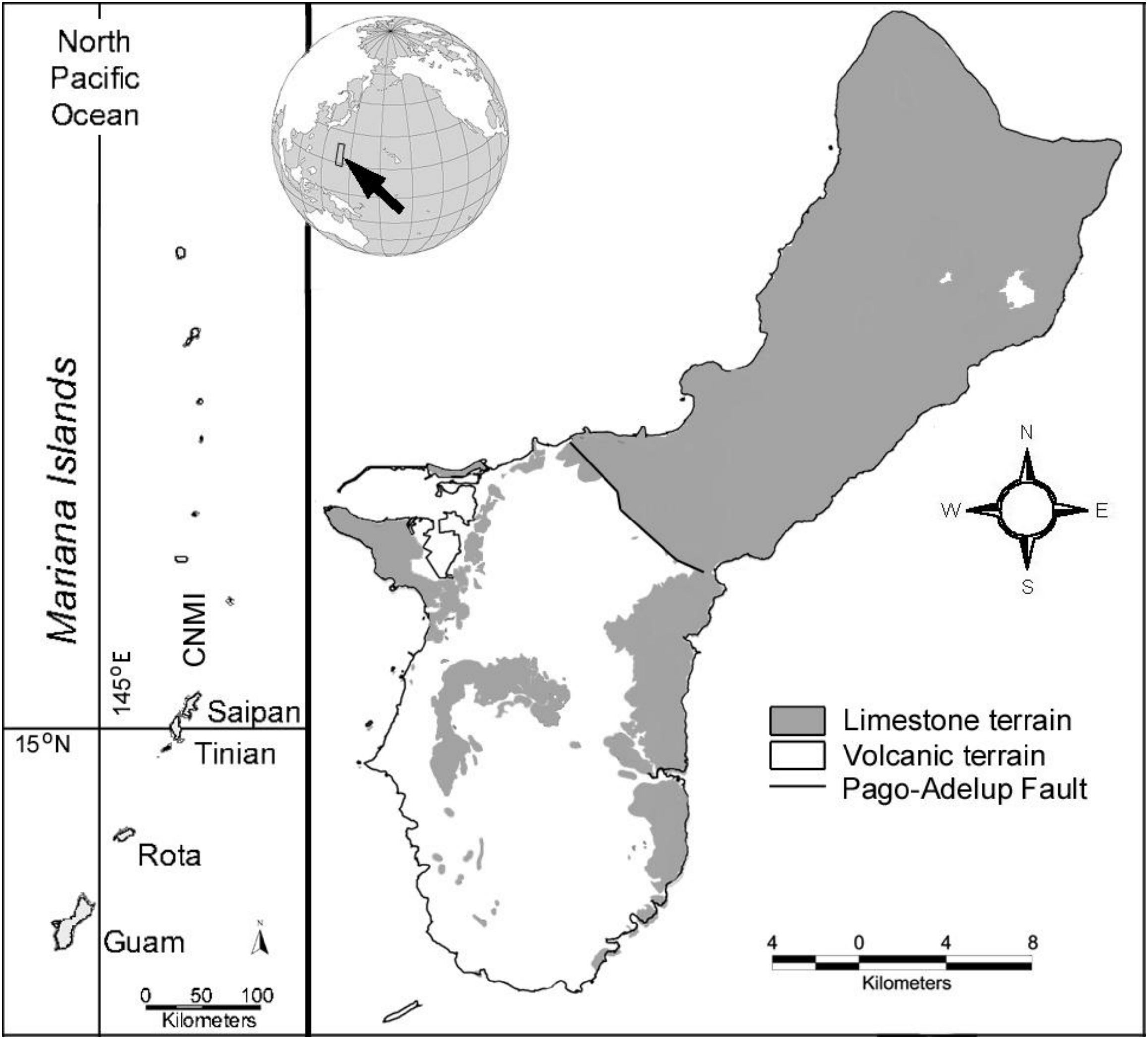


**Fig. 1. Location of Guam and the extent of its limestone terrain over the two provinces separated by the Pago-Adelup Fault.**
